# Dopamine and its receptor *DcDop2* are involved in the coevolution between ‘*Candidatus* Liberibacter asiaticus’ and *Diaphorina citri*

**DOI:** 10.1101/2025.10.10.681724

**Authors:** Xiaoge Nian, Jiayun Li, Jilei Huang, Weiwei Yuan, Paul Holford, George Andrew Charles Beattie, Jielan He, Yijing Cen, Yurong He, Songdou Zhang

## Abstract

‘*Candidatus* Liberibacter asiaticus’ (CLas), the causal agent of citrus huanglongbing, is transmitted by the Asian citrus psyllid *Diaphorina citri*. While *C*Las-positive (*C*Las+) females exhibit increased fecundity and metabolic demands, their neuroendocrine regulation mechanisms remain unclear. We propose *C*Las manipulates dopamine (DA) signaling to enhance psyllid fecundity and *C*Las proliferation. Metabolomics revealed elevated DA in *C*Las+ females. Silencing DA synthesis genes and receptor *DcDop2* via RNAi reduced lipid reserves, fecundity, and ovarian *C*Las titers. Through combined *in vivo* and *in vitro* experiments, we demonstrated that the microRNA miR-31a suppresses *DcDop2* expression by binding to its 3’ untranslated region. Overexpression of miR-31a resulted in decreased *DcDop2* expression and *C*Las titers in the ovaries, eliciting phenotypic defects akin to *DcDop2* knockdown. Furthermore, *DcDop2* knockdown and miR-31a overexpression reduced juvenile hormone (JH) levels and adipokinetic hormone (AKH) signaling in fat bodies and ovaries. Consequently, *C*Las regulates the DA-*DcDop2* signaling axis to improve *D. citri* lipid metabolism and fecundity, while simultaneously promoting its replication. These findings reveal a coevolution between *C*Las proliferation and ovarian development in the insect host. This discovery enhances our understanding of the molecular interplay between plant pathogens and vector insects and offers novel targets and strategies for HLB field management.

## Introduction

Plant pathogens can significantly reduce the yields and quality of agricultural and horticultural crops, and many rely on arthropod vectors for their transmission from host to host (Bourne et al., 2024). Over recent decades, extensive research has focused on the intricate relationships that can occur between vectors and pathogens (Mao et al., 2019; Berasategui et al., 2022; Ray and Casteel, 2022). Exploring such interactions between the pathogens and vectors stands at the forefront of vector biology and ecology, offering a foundational framework for the efficacious prevention and management of vector-borne plant diseases. Nonetheless, investigations focused on the mechanisms governing these interactions have predominantly centered on vector-virus or vector-fungus dynamics, with comparatively less attention devoted to vector-bacteria interactions (Mao et al., 2019; Ray and Casteel, 2022).

The most serious impediment to global citrus production is the severe Asiatic form of a disease known as huánglóngbìng (HLB: yellow shoot disease) (Beattie, 2020a; Beattie, 2020b; Fuentes et al., 2018; Gottwald et al., 2010; Leong et al., 2022; Pérez-Hedo et al., 2024; Wang et al., 2017). It is associated with ‘*Candidatus* Liberibacter asiaticus’ (*C*Las: α-Proteobacteria) a phloem-limited bacterium that is normally transmitted between rutaceous hosts by the Asiatic citrus psyllid (ACP), *Diaphorina citri* Kuwayama (Hemiptera: Psyllidae). The disease has severely affected citrus cultivation in Asia, initially in South Asia since late 1800s, and in the Americas since 2004. *C*Las and ACP both originated in South Asia (Leong et al., 2022). The original host of the pathogen is not known. The original host of ACP was probably a citrus relative, possibly curry leaf (*Bergera koenigii* L.) or *Murraya intermedia* (M. Roem.) Mabb (Beattie, 2020a). Koch’s postulates confirming *C*Las as the cause of the disease have not fully demonstrated; partial fulfillment was reported by Zheng et al. (2024) and there is general agreement in the research community that the disease is caused by *C*Las. Use of insecticides is the most common strategy used for minimizing populations of *D. citri* to limit impacts of the disease (Gottwald et al., 2010; Bové, 2014). However, experience in Asia (Beattie, 2020b; Leong et al., 2022), the United States of America (Graham et al., 2020; Li et al., 2020), and recent media reports from Brazil, indicate that it is not possible to rely on use insecticides to prevent spread of *C*Las.

Recent studies have indicated that *C*Las-positive *D. citri* exhibit higher fecundity in comparison to their *C*Las-negative counterparts (Pelz-Stelinski and Killiny, 2016; Ren et al., 2016; Wu et al., 2018). Nian et al. (2024) reported that infection with *C*Las enhances the fecundity of *D. citri* and then aids the pathogen proliferation; a win-win strategy. Furthermore, Li et al. (2024) demonstrated that adipokinetic hormone (AKH)/ AKH receptor (AKHR) signaling influences lipid metabolism and boosts the fecundity of *D. citri* induced by *C*Las. Reproduction during the adult life stage of female insects is a metabolically demanding process intricately governed by the neurohormone system; a delicate balance exists between neural control and lipid storage and utilization throughout the reproductive phase (Siju et al., 2021). Nevertheless, the precise mechanism about how *D. citri* maintains this balance following *C*Las infection, involving neurohormone regulation, lipid metabolism, and heightened fecundity, still lacks comprehensive elucidation.

Dopamine (DA), a conserved and potent catecholamine neurotransmitter found in both vertebrates and invertebrates, plays crucial roles in modulating various behaviors such as locomotor activity, sexual behavior, development, reproduction, and endocrine functions (Ma et al., 2021; Sasaki et al., 2017; Shen et al., 2020; Morigami and Sasaki, 2024). The fundamental steps involved in dopaminergic neurotransmission are evolutionarily conserved between flies and humans. In *Drosophila melanogaster* Meigen (Diptera: Drosophilidae), tyrosine hydroxylase (TH), the rate-limiting enzyme in dopamine biosynthesis and encoded by the *pale* gene, catalyzes the conversion of tyrosine to l-3,4-dihydroxyphenylalanine (L-DOPA), an intermediate subsequently transformed into DA via decarboxylation by dopa decarboxylase (DDC). The vesicular amine transporter (VAT) governs the synaptic release of DA, activating DA receptors to initiate downstream signal transduction pathways. Vertebrates possess five DA receptor subtypes categorized into D1-like (D1 and D5) and D2-like (D2, D3, D4) receptor groups (Xu et al., 2017). In contrast, invertebrates, particularly insects, have four DA receptor subtypes: D1-like DA receptor (Dop1), invertebrate-type DA receptor (Dop2), D2-like DA receptor (Dop3), and DopEcR. Dopamine and its receptors play crucial roles in insect reproductive processes (Sasaki et al., 2017). In male *D. melanogaster*, brain DA levels influence sexual orientation, with both excessively high and low DA levels triggering same-sex courtship behavior among males (Liu et al., 2008; Chen et al., 2012). Mutations in DA receptors (Dop1, Dop2, Dop3, and DopEcR) in *D. melanogaster* result in reduced mating time (Crickmore and Vosshall, 2013). In *Tribolium castaneum* (Herbst) (Coleoptera: Tenebrionidae), disruption of Dop3 inhibits yolk protein uptake and ovarian development (Bai and Palli, 2016). Whether DA signaling involves in the heightened fecundity of *C*Las-positive *D. citri* currently remains unknown.

While extensive research has focused on the DA signaling pathway, the post-transcriptional regulation of this pathway by microRNAs (miRNAs) remains relative understudied. miRNAs, small non-coding RNAs approximately 22 nucleotides in length, bind to the 3’-untranslated region (UTR) of target messenger RNAs (mRNAs), leading to either translation repression or mRNA degradation (Bartel, 2018). These miRNAs play pivotal roles in modulating cellular processes at the post-transcriptional level across various biological contexts (Lucas et al., 2015; Chen et al., 2020; Song et al., 2020). In the migratory locust, *Locusta migratoria* (L.) (Orthoptera: Acrididae), miRNA-133 targets two key genes, *henna* and *pale*, within the DA synthesis pathway, resulting in the downregulation of their expression and subsequent inhibition of DA synthesis. This inhibition leads to the transition of locusts from gregarious to solitary phases (Yang et al., 2014). Additionally, Dop1 can mediate locust olfactory attraction behavior by inhibiting the expression of miR-9a (Guo et al., 2018). In *Helicoverpa armigera* (Hübner) (Lepidoptera: Noctuidae), miR-14 and miR-2766 are involved in regulating larval metamorphosis by targeting the expression of *HaTH* that encodes tyrosine hydroxylase (Shen et al., 2022). Recent findings by Liu et al. (2023) demonstrated that the miRNA, let-7, increases the sensitivity of European honeybee, *Apis mellifera* L. (Hymenoptera: Apidae), to sucrose by targeting the DA receptor, *AmDop2*. Currently, there is a dearth of studies elucidating the role of miRNAs in regulating genes within the DA signaling pathway of citrus psyllids.

In a previous study, we observed that *C*Las regulates the juvenile hormone (JH) signaling pathway and host miR-275, which targets the *vitellogenin receptor* (*DcVgR*), to improve *D. citri* fecundity while simultaneously advance *C*Las replication (Nian et al., 2024). This suggests a coevolution between *C*Las and its insect vector. Subsequent research showed that *C*Las manipulates AKH/AKHR-JH signaling to enhance *D. citri* lipid metabolism and fecundity, meanwhile facilitating its own replication. As of now, there is a lack of reports detailing the involvement of miRNAs targets DA signaling genes in *D. citri*-*C*Las interactions that cause the increased fecundity of *C*Las-infected *D. citri* females. Therefore, in this study, we utilize the *D. citri*-*C*Las interaction system as a model to explore the molecular mechanisms underpinning the influence of DA signaling on lipid metabolism and fecundity of *C*Las-positive *D. citri*. Our research aims to enhance our understanding of interactions between insect vectors and pathogens, with the potential to reveal innovative strategies for managing *D. citri* and HLB.

## Results

### *C*Las infection elevates dopamine levels and the expression of dopamine biosynthesis genes in female adult *D. citri*

To investigate the potential association between neurotransmitters and increased fecundity of *C*Las-positive (*C*Las+), female *D. citri*, we constructed six libraries using samples from *C*Las+ and *C*Las-negative (*C*Las-) individuals for metabolomic analyses of neurotransmitter contents. The analyses revealed alterations in 23 neurotransmitters: 6 were upregulated, 7 were downregulated, 5 remained unchanged, and 5 were undetectable. Notably, among the upregulated neurotransmitters, levels of DA and L-DOPA were significantly higher in *C*Las+ females compared to their *C*Las− counterparts (Figure 1A). Further investigations confirmed the involvement of DA in the *D. citri*-*C*Las interaction by assessing DA levels in *C*Las− and *C*Las+ females. Remarkably, DA levels showed a significant increase during the ovarian development stages 5, 9, and 13 day after emergence (DAE) in *C*Las+ females compared to *C*Las− females (Figure 1B). In insects, DA production is intricately linked to the reproductive process, involving the conversion of phenylalanine (Phe) to tyrosine (Tyr), tyrosine hydroxylation to generate L-DOPA, decarboxylation of L-DOPA to form DA, and subsequent release of DA to target tissues (Figure 1C). To assess the expression profiles of genes associated with DA biosynthesis and release, we analyzed the expression of *DcHenna1* and *DcHenna2* (encoding phenylalanine hydroxylase 1 and 2), *DcTh* (encoding tyrosine hydroxylase), *DcDdc* (encoding dopa decarboxylase), and *DcVat1 and DcVat2* (encoding vesicular amine transporter 1 and 2). These genes were chosen based on transcriptomic sequencing conducted in our laboratory. Using qPCR data, we created the heat maps illustrating the expression levels of these genes among various tissues. Notably, compared to *C*Las− females, elevated mRNA levels of *DcHenna1*, *DcTh*, *DcDdc*, and *DcVat1* were detected in the ovaries, whole body and head of *C*Las+ females (Figure 1D and S1). These findings imply a possible role of the DA signaling pathway in modulating the increased fecundity seen in *C*Las+ female adults.

**Figure 1.**
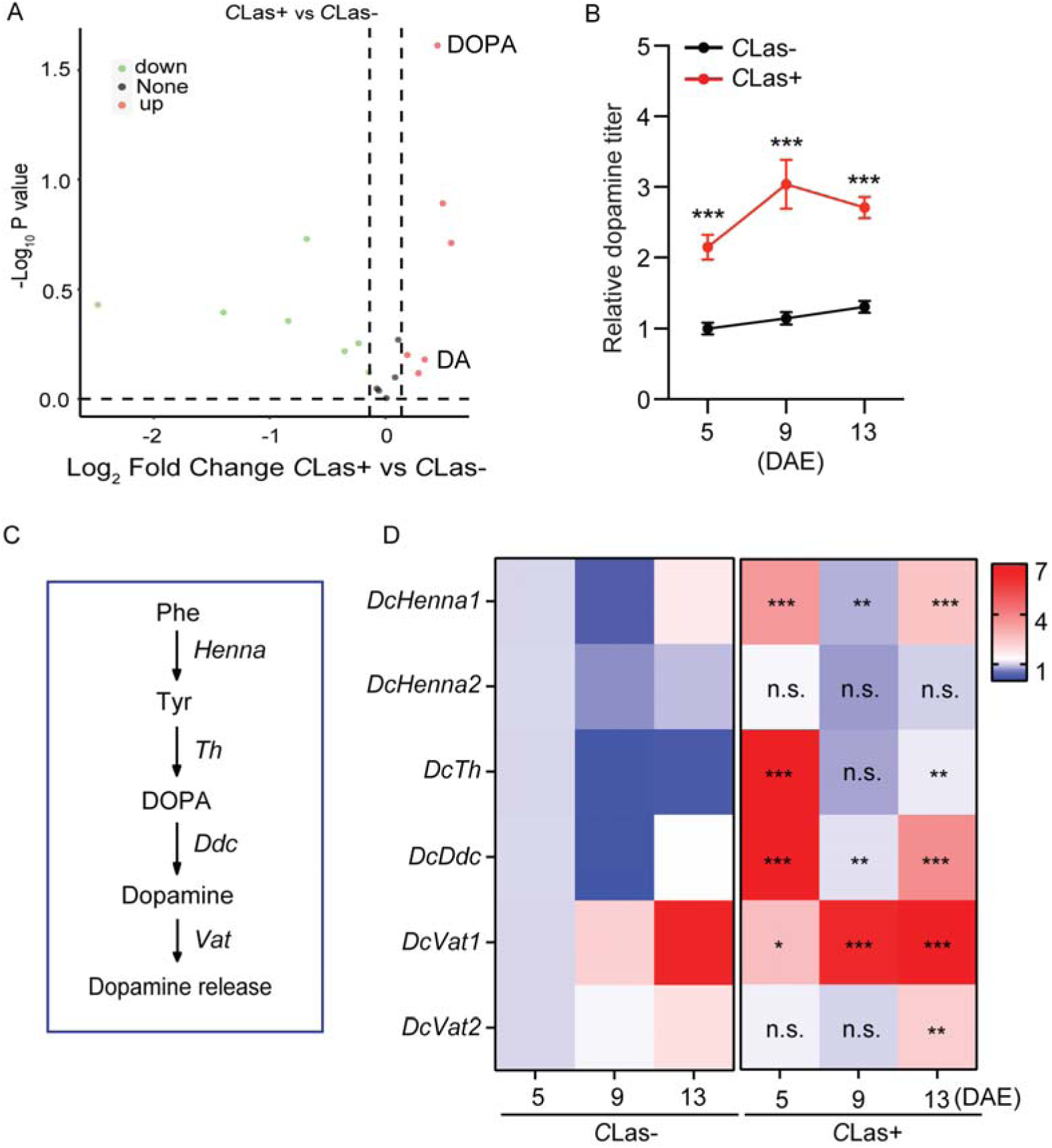
Effects of *C*Las on dopamine levels and the expression of DA biosynthesis genes in *D. citri.* (A) A volcano plot highlighting differentially-expressed neurotransmitters in *C*Las+ and *C*Las− psyllid ovaries, with six being upregulated (red dots) and seven downregulated (green dots). (B) Comparison of dopamine levels in the ovaries of *C*Las+ and *C*Las− females 5, 9, 13 days after emergence (DAE). (C) Schematic representation of dopamine biosynthesis and release pathway. (D) A heatmap indicating the temporal expression patterns of six dopamine metabolic pathway genes. Data in 1B and 1D are presented as means ± SEM with three independent biological replicates, each with three technical replicates. Significant differences between *C*Las− and *C*Las+ psyllids were determined using Student’s *t*-tests (\**p* <0.05, \*\**p* < 0.01, \*\*\**p* < 0.001).

### Pivotal genes involved in DA biosynthesis regulate the changes in metabolism and fecundity in *D. citri* mediated by *C*Las

To investigate the role of DA in the *D. citri*-*C*Las interaction within the ovaries, we utilized RNAi to target crucial genes, specifically *DcHenna1*, *DcTh*, *DcDdc*, and *DcVat1*. Following exposure to ds*RNA*, the mRNA levels of these genes in the ovaries of *C*Las− and *C*Las+ females were significantly reduced compared to those treated with ds*GFP* (Figure S2). Knockdown of these four genes led to a marked decrease in the dopamine titer (Figures 2A), as well as triacylglycerol (TAG) contents, glycogen levels, and lipid droplet sizes in both *C*Las-and *C*Las+ females (Figures 2B-D). Additionally, the reduced expression of these critical genes disrupted ovarian development, extended the preoviposition periods, shortened the oviposition periods, and reduced the fecundity compared to ds*GFP* treatments in both *C*Las-and *C*Las+ females (Figure 2E-H). Furthermore, the *C*Las signal intensity and relative *C*Las titers in ovaries were notably diminished after gene knockdown (Figures 2I-J). Altogether, the DA signaling pathway acts as a crucial regulator of energy metabolism and reproductive enhancement triggered by *C*Las in infected *D. citri* females.

**Figure 2.**
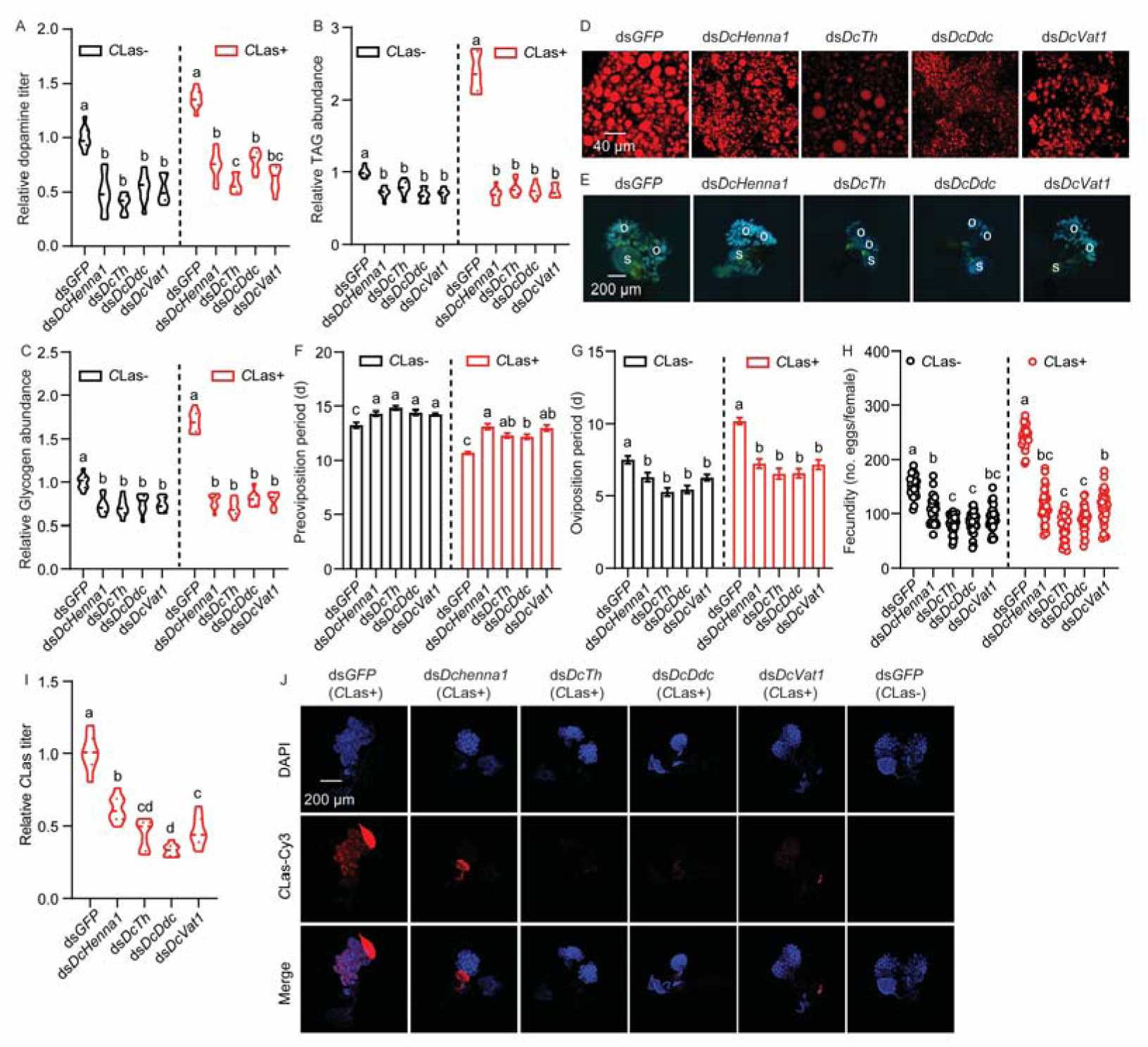
Effects of RNAi-mediated silencing of *DcHenna1*, *DcTh*, *DcDdc*, and *DcVat1* on energy metabolism and fecundity of *C*Las+ females. (A) Dopamine titer in the *C*Las− and *C*Las+ females post treatment for 48 h with ds*DcHenna1*, *dsDcTh*, *dsDcDdc*, or *dsDcVat1*. (B-C) TAG and glycogen levels in the fat bodies of *C*Las− and *C*Las+ females post treatment for 48 h with ds*DcHenna1*, *dsDcTh*, *dsDcDdc*, or *dsDcVat1*. (D) Lipid droplets stained with Nile red in fat bodies dissected from *C*Las+ females treated with ds*DcHenna1*, ds*DcTh*, ds*DcDdc*, or ds*DcVat1*. Scale bar = 40 μm. (E) Ovary phenotypes in *C*Las+ females post *DcHenna1*, *DcTh*, *DcDdc*, and *DcVat1* treatment. Scale bar = 200 μm. o: ovary, s: spermathecae. (F-H) Comparison of preoviposition period, oviposition period, and fecundity between *C*Las− and *C*Las+ females treated with ds*DcHenna1*, ds*DcTh*, ds*DcDdc*, or ds*DcVat1*. (I) *C*Las titer in ovaries of *C*Las+ females at 7 DAE treated with ds*DcHenna1*, ds*DcTh*, ds*DcDdc*, or ds*DcVat1*. (J) Representative confocal images of *C*Las in reproductive systems of *C*Las+ females treated with ds*DcHenna1*, ds*DcTh*, ds*DcDdc*, or ds*DcVat1*. Scale bar = 200 μm. DAPI: cell nuclei were stained with DAPI and visualized in blue. *C*Las-Cy3: *C*Las signal visualized in red by staining with Cy3. Merge: merged imaging of co-localization of cell nuclei and *C*Las. Results in 2A-2C and 2I were displayed as means ± SEM with three independent biological replicates, each with three technical replicates. Data in 2F-2H were shown as means ± SEM with thirty independent biological replicates. For A-C, and F-I, significant differences among different treatments are denoted by lowercase letters based on one-way ANOVA followed by the Tukey’s HSD tests at *p* < 0.05.

### *DcDop2* acts as a DA receptor and participates in *D. citri*-*C*Las mutualism

To identify the DA receptor, fragments of three putative DA receptors in *D. citri* were obtained from the National Center for Biotechnology Information (NCBI, https://www.ncbi.nlm.nih.gov/, 1988) database, and designated as *DcDop1* (XM_017446452.2), *DcDop2* (XM_026822145.1), and *DcDop3* (XM_008472271.3). The full coding sequence of *DcDop1* consist of 1281 nucleotides encoding 426 amino acids, *DcDop2* comprises 2049 nucleotides encoding 682 amino acids, and *DcDop3* comprises 1560 nucleotides encoding 519 amino acids. To elucidate the evolutionary relationships between *D. citri* DA receptors and homologues from other insect species, a phylogenetic tree was constructed, revealing a close clustering of *DcDop1*, *DcDop2*, and *DcDop3* with their orthologous receptors (Figure S3). Due to the presence of three DA receptors, the response of each to elevated DA levels induced by *C*Las infection was assessed using qRT-PCR. The results showed that only *DcDop2* responded with an increase in transcription during the evaluation period, with higher levels in *C*Las+ females compared to CLas− females (Figure 3A); minimal changes in the expression *DcDop1* and *DcDop3* were observed (Figure S4). Furthermore, tissue-specific expression patterns revealed that *DcDop2* exhibited the highest expression in the midgut, followed by the head, fat bodies, and ovaries (Figure S5). To validate the up-regulation of *DcDop2* by DA, a cell-based calcium mobilization assay was performed; the assay showed significant activation of *DcDop2* by DA in a dose-dependent manner, with an EC_50_ value of 4.05 × 10^-6^ mM; no response was observed in an empty vector when challenged with DA (Figure S6).

**Figure 3.**
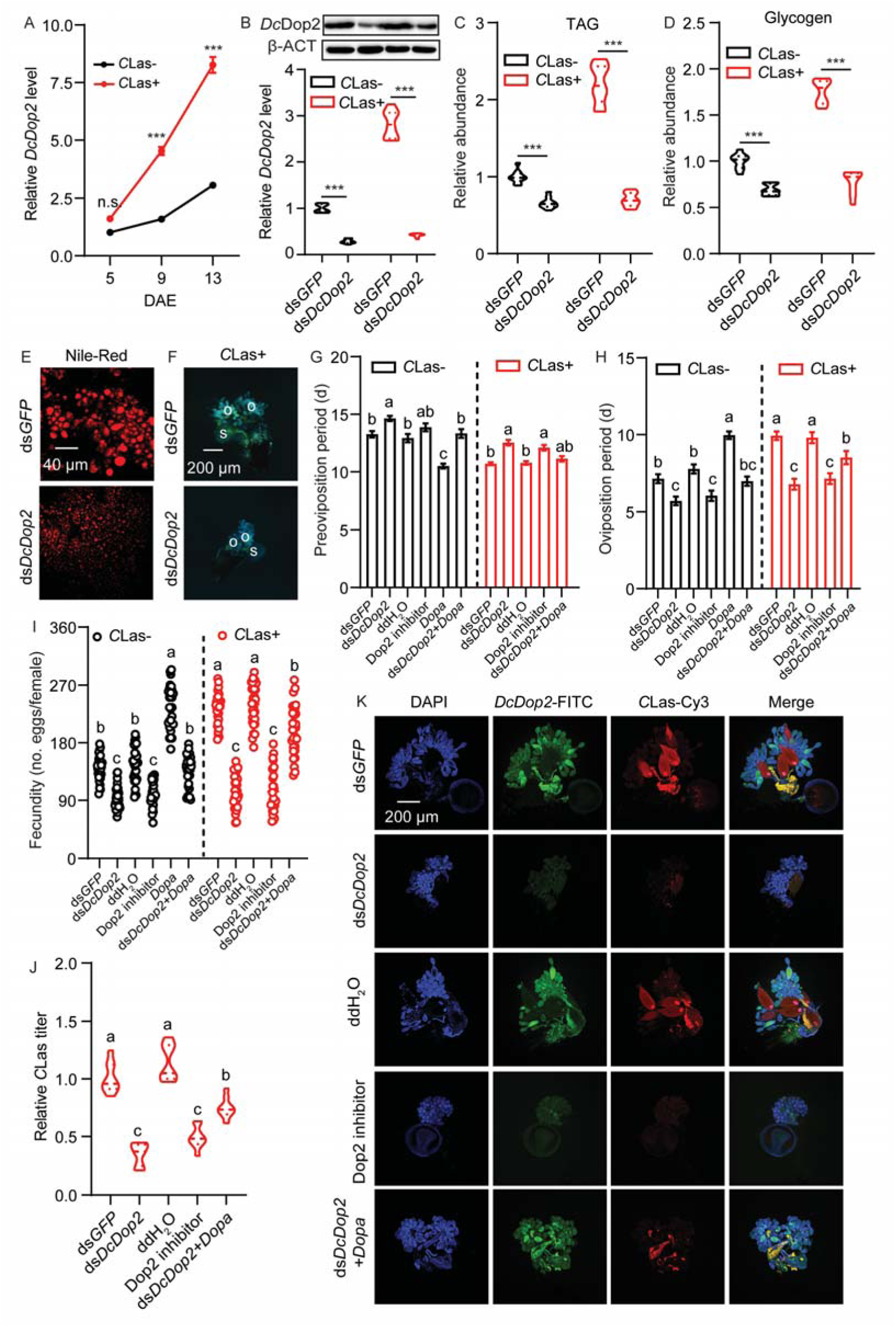
Dopamine concentrations mediated by *DcDop2* are involved in the coevolution between *C*Las and *D. citri* resulting in increased fecundity. (A) Temporal expression patterns of *DcDop2* between the ovaries of *C*Las− and *C*Las+ females. (B) Effects of RNAi on *DcDop2* expression at protein and mRNA levels in *C*Las− and *C*Las+ females treated with ds*DcDop2* for 48 h. (C-D) Comparison of TAG and glycogen levels in fat bodies of *C*Las− and *C*Las+ females treated with ds*DcDop2* for 48 h. (E) Lipid droplets stained with Nile red in fat bodies dissected from *C*Las+ females treated with ds*DcDop2* for 48 h. Scale bar = 40 μm. (F) Ovary phenotypes in ds*DcDop2*-treated *C*Las+ females at 48 h. Scale bar = 200 μm. o: ovary, s: spermathecae. (G-I) Comparison of preoviposition period, oviposition period, and fecundity of *C*Las− and *C*Las+ adults treated with ds*DcDop2*, Dop2 inhibitor (Pimozide, HY-12987, MedChemExpress), and dopamine rescue. (J) *C*Las titer in ovaries of *C*Las+ females at 7 DAE treated with ds*DcDop2*, Dop2 inhibitor (Pimozide, HY-12987, MedChemExpress), and DA rescue. (K) Representative confocal images of the reproductive system of *C*Las+ females treated with ds*DcDop2*, Dop2 inhibitor (Pimozide, HY-12987, MedChemExpress), and dopamine rescue. Scale bar = 200 μm. DAPI: cell nuclei were stained with DAPI and visualized in blue. *DcDop2*-FITC: *DcDop2* signal visualized in green by staining with FITC. *C*Las-Cy3: *C*Las signal visualized in red by staining with Cy3. Merge: merged imaging of co-localization of cell nuclei, *DcDop2* and *C*Las. Results in 3A-3D and 3J were displayed as means ± SEM with three independent biological replicates, each with three technical replicates. Data in 3G-3I were shown as means ± SEM with thirty independent biological replicates. Significant differences between treatments and controls are indicated by asterisks in A-D (Student’ s *t*-test, \**p* < 0.05, \*\**p* < 0.01, \*\*\**p* < 0.001). For G-J, significant differences among different treatments are indicated by lowercase letters based on one-way ANOVA followed by Tukey’s HS tests at *p* < 0.05.

To explore the influence of *DcDop2* on energy metabolism and the reproductive alterations induced by *C*Las, RNAi was utilized to suppress *DcDop2* expression. Psyllids fed with ds*DcDop2* displayed a substantial decrease in *DcDop2* at protein and mRNA levels in the ovaries of both CLas− and CLas+ females by nearly 80%, respectively, compared to those fed with ds*GFP* (Figure 3B). Silencing of *DcDop2* led to significant reductions in TAG and glycogen levels, along with a notable decrease in lipid droplet sizes (Figure 3C-E). Additionally, *DcDop2* RNAi disrupted ovarian development (Figure 3F), extended the preoviposition period (from 13.3 to 14.6 d in *C*Las− females and from 10.7 to 12.5 d in *C*Las+ females) (Figure 3G), shortened the oviposition period (from 7.1 to 5.7 d in *C*Las− females and from 9.9 to 6.8 d in *C*Las+ females) (Figure 3H), and reduced fecundity (from 140.6 to 94.0 eggs per female in *C*Las− females and from 233.3 to 104.5 eggs per female in *C*Las+ females) (Figure 3I). When an inhibitor was used to decrease the mRNA expression of *DcDop2*, it resulted in phenotype abnormalities similar to those observed with RNAi-mediated *DcDop2* knockdown. These effects included an extended preoviposition period, a shortened oviposition period, decreased fecundity, as well as reductions in *C*Las titer and signal intensity. Interestingly, DA treatment was able to rescue the defective phenotype arising from RNAi-mediated *DcDop2* knockdown in *C*Las− females and *C*Las+ females. Furthermore, the *C*Las signal intensity and relative *C*Las titer in the ovaries of females with *DcDop2* knockdown were significantly diminished in the follicle cells and oviduct (Figures 3J-K). These findings suggest that *DcDop2* responds to DA and plays a role in the *C*Las-induced changes in energy metabolism and reproduction in *D. citri*.

### miR-31a directly targets *DcDop2* by binding to its 3’-UTR

Utilizing miRanda and RNAhybrid software, we predicted potential miRNAs that could target *DcDop2* from our small, *D. citri* RNA libraries. Three putative miRNAs, namely miR-31a, miR-275, and miR-15 (Figure 4A) were identified and predicted to bind to 3’-UTR sequences. To assess the binding activity of these miRNAs with *DcDop2 in vitro*, dual-luciferase reporter assays were conducted. Notably, co-transfection of miR-31a agomir (agomir-31a) with the recombinant plasmid containing the full 3’-UTR sequence of *DcDop2* led to a significant decrease in luciferase activity compared to the controls, while the activities in the miR-275 or miR-15 treatment groups remained unchanged (Figure 4B). Moreover, the reporter activity was restored upon mutation of the binding sites of miR-31a in the3’-UTR of *DcDop2* (Figure 4C). To confirm the specific targeting of *DcDop2* by miR-31a, a series of experiments were conducted. Firstly, tissue expression patterns revealed high expression levels of miR-31a in the head, followed by the fat bodies, ovaries, and midgut (Figure 4D). Secondly, miR-31a displayed an expression pattern opposite to that of *DcDop2*, with transcript levels decreasing over the evaluation period (Figure 4E). Thirdly, treatment with agomir-31a significantly increased miR-31a expression in the ovaries of both *C*Las− and *C*Las+ females (Figure S7). Correspondingly, the mRNA levels of *DcDop2* either increased or decreased after treatments with either antagomir-31a or agomir-31a (Figure 4F), separately. Fourthly, an RNA immunoprecipitation (RIP) assay demonstrated a significant enrichment of *DcDop2* mRNA in the anti-AGO-immunoprecipitated RNAs from the ovaries of female psyllids fed with agomir-31a compared to the control samples (Figure 4G). Collectively, these findings strongly support the conclusion that *DcDop2* is a direct target of miR-31a.

**Figure 4.**
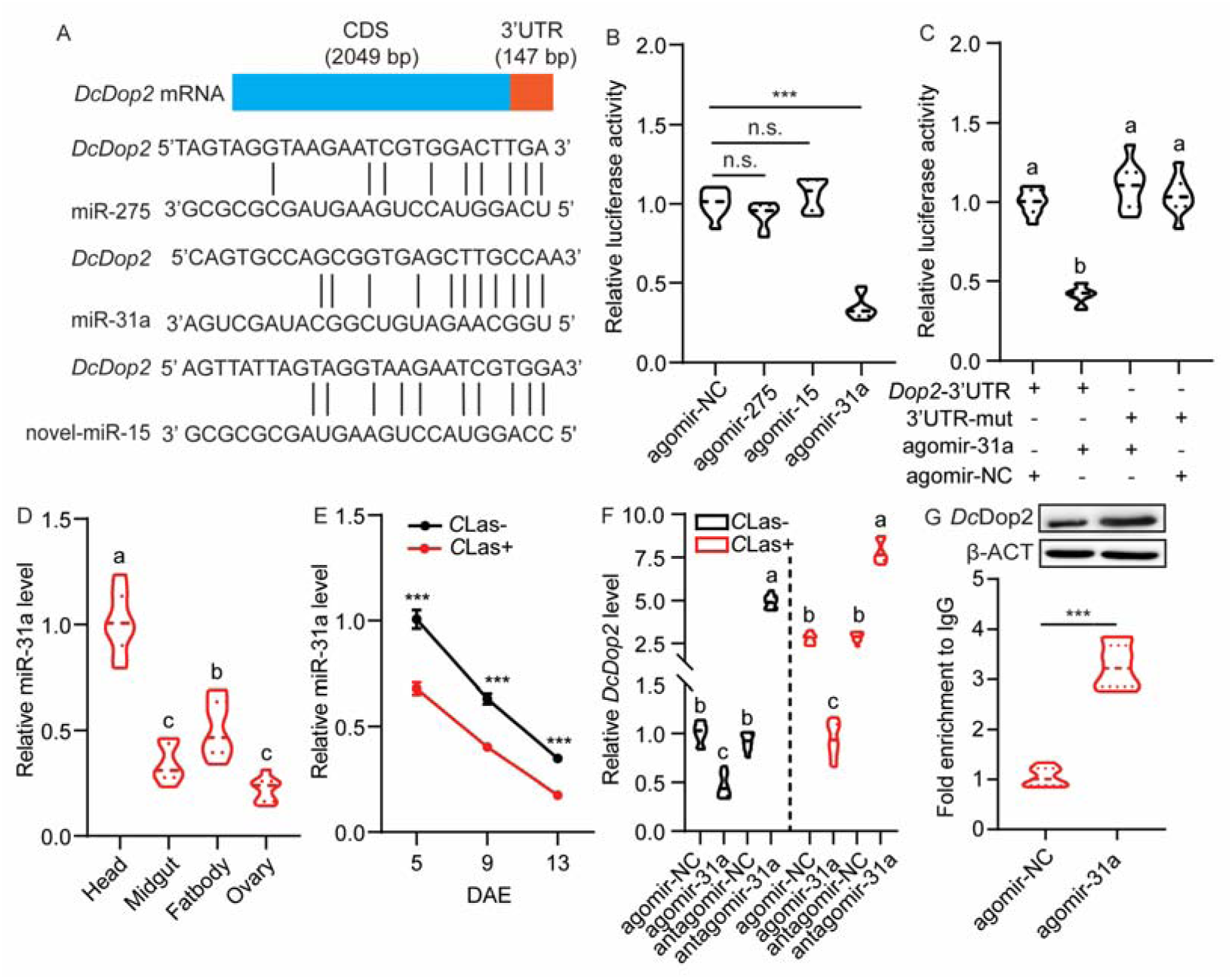
Identification and validation of the targeting of *DcDop2* by miR-31a. (A) Prediction of potential miRNA binding sites in the 3’-UTR of *DcDop2* using miRanda and RNAhybrid. (B) Assessment of miRNA-mediated regulation through dual-luciferase reporter assays in HEK293T cells co-transfected with miRNA agomir and recombinant pmirGLO vectors containing binding sites for miR-275, novel-miR-15 and miR-31a in the 3’ UTR of *DcDop2.* (C) Dual-luciferase reporter assays in HEK293T cells co-transfected with miR-31a agomir and recombinant pmirGLO vectors containing will-type or mutated *DcDop2-*3’UTR. (D) Tissue-specific expression of miR-31a in *C*Las+ female adults 7 DAE in the head, ovary, fat body, and midgut. (E) Comparison of temporal expression patterns of miR-31a in the ovaries of *C*Las− and *C*Las+ females. (F) Impact of miR-31a agomir and antagomir treatments on *DcDop2* mRNA level in the ovaries of *C*Las− and *C*Las+ psyllids after 48 h. (G) *In vivo* assessment of miR-31a targeting *DcDop2* through RNA immunoprecipitation assay. Data in 4B-4G are presented as means ± SEM with three independent biological replicates, each with three technical replicates. For figures C, D, and F, significant differences among the different treatments are indicated by lowercase letters based on one-way ANOVA followed by Tukey’s Honest Significant Difference tests at *p* < 0.05. Significant differences between treatments and controls are denoted by asterisks in B and E-G (Student’s *t*-test, \*\**p* < 0.01, \*\*\**p* < 0.001).

### miR-31a participates in *D. citri*-*C*Las mutualism within ovaries

To investigate the influence of miR-31a on the *D. citri* and *C*Las interaction, both *C*Las-and *C*Las+ females were administered either agomir-NC or agomir-31a. Upon agomir-31a treatment, both populations displayed a significant decrease in TAG and glycogen contents (Figures 5A-B). The lipid droplets in the fat bodies of *C*Las+ females appeared smaller compared to those in the control group (Figure 5C). Furthermore, treatment with agomir-31a led to impaired ovarian development in *C*Las+ females (Figure 5D). Additionally, exposure of both *C*Las− and *C*Las+ females to agomir-31a resulted in a shortened oviposition period, a significantly prolonged preoviposition period, and a marked decrease in fecundity in comparison to their respective control groups (Figures 5E-G). These observed phenotypes closely resembled those induced by ds*DcDop2* treatment. Moreover, a noticeable reduction in *C*Las signal and relative titers was observed in *C*Las+ ovaries (Figures 5H-I). In conclusion, these results suggest that miR-31a suppresses *DcDop2* expression and plays a critical role in the coevolution between *D. citri* and *C*Las within the ovaries.

**Figure 5.**
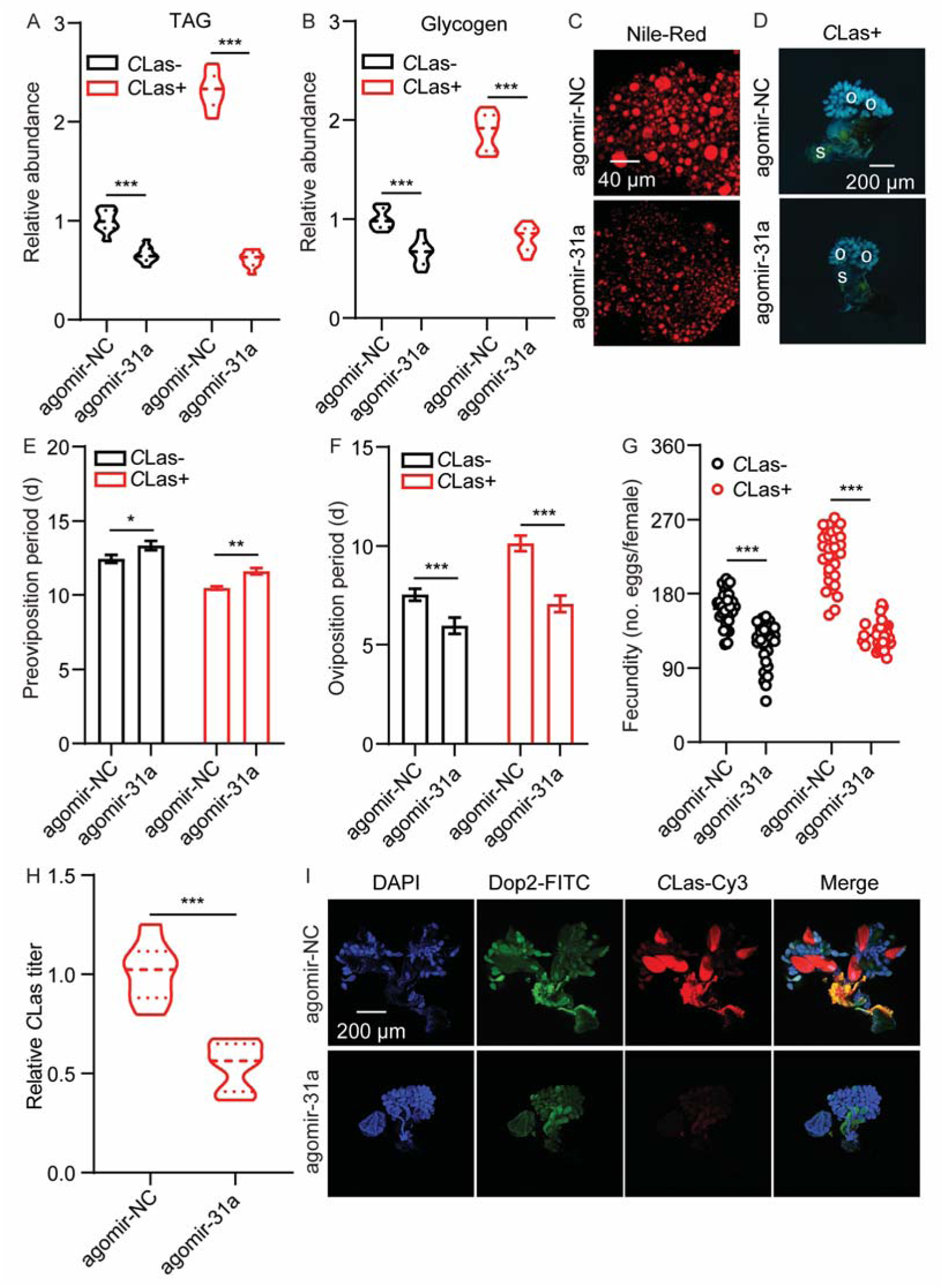
Involvement of miR-31a in the coevolution between *D. citri* and *C*Las. (A-B) Comparative analysis of TAG and glycogen levels in the fat bodies of *C*Las− and *C*Las+ females following agomir-31a treatment for 48 h. (C) Visualization of lipid droplets stained with Nile red in fat bodies extracted from *C*Las+ females treated with agomir-31a for 48 h. Scale bar = 40 μm. (D) Ovarian phenotypes of *C*Las+ female treated with agomir-31a for 48 h. Scale bar = 200 μm. o: ovary, s: spermathecae. (E-G) Comparison of the preoviposition period, oviposition period, and fecundity between *C*Las− and *C*Las+ adult females treated with agomir-31a. (H) Quantification of *C*Las titer in the ovaries of *C*Las+ females at 7 DAE treated with agomir-31a for 48 h. (I) Representative confocal images showing *DcDop2* and *C*Las in the reproductive system of *C*Las+ females treated with agomir-31a for 48 h. Scale bar = 200 μm. The signals of DAPI, *DcDop2*-FITC, and *C*Las-Cy3 are consistent with described in Figure 2. Results in 5A-5B and 5H were displayed as means ± SEM with three independent biological replicates, each with three technical replicates. Data in 5E-5G were shown as means ± SEM with thirty independent biological replicates. Significant differences between treatments and controls indicated by asterisks (Student’s *t*-test, \*\*\**p* < 0.001).

### The modulation of AKH and JH signaling pathway by the DA pathway contributes to enhanced fecundity of *C*Las+ female *D. citri*

To assess the potential regulatory impact of DA signaling pathway on the AKH and JH signaling pathways, we examined the relative JH titers and the expression levels of key genes involved in these pathways in *DcDop2*-deficient *C*Las+ females. Following ds*DcDop2* treatment, a significant decrease in the relative JH titer in the abdomen was observed after 48 h in *C*Las+ females compared to ds*GFP* control females (Figure 6A). Moreover, the expression levels of key genes the AKH signaling (*DcAKH* and receptor gene *DcAKHR*) and JH signaling (receptor gene *DcMet* and the downstream transcription factor *DcKr-h1*) were notably reduced in the fat bodies and ovaries after 48 h of ds*DcDop2* treatment (Figures 6B-C). Additionally, the expression levels of key downstream genes of the JH signaling pathway involved in ovarian development, such as *DcVg1-like*, *DcVgA1-like*, and *DcVgR*, were also lower in *DcDop2*-deficient *C*Las+ females following dsRNA feeding for 48 h, compared to controls (Figures 6B-6C). Interestingly, JHA and AKH was able to partially rescue the defective phenotype arising from RNAi-mediated *DcDop2* knockdown in *C*Las+ females (Figures 6D-E), which means JH and AKH function as the downstream of DA signaling. Similar trends were observed in *C*Las+ females treated with agomir-31a (Figures 6F-I). These findings suggest that the DA-*DcDOP2*-miR-31a signaling axis as a newly identified upstream regulator of the existing AKH and JH signaling pathways, thereby contributing to the increased fecundity observed in *C*Las+ *D. citri* females.

**Figure 6.**
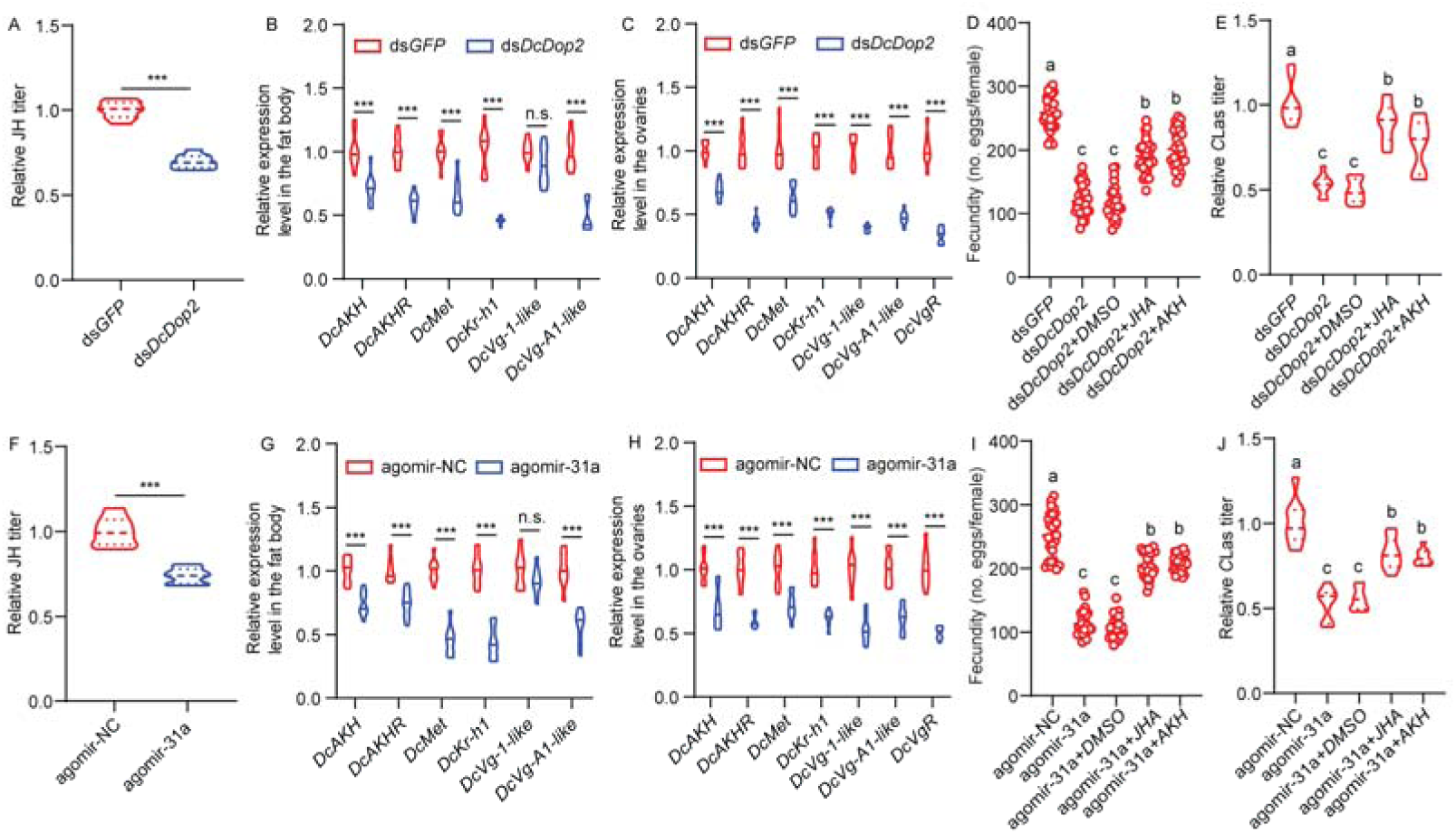
The modulation of the AKH and JH signaling pathway by the DA pathway enhances fecundity in *C*Las+ *D. citri.* (A) JH titer in the abdomen of *C*Las+ females treated with ds*DcDop2* for 48 h. (B) Influence of ds*DcDop2* treatment on mRNA levels of key genes in the AKH and JH signaling pathway in fat bodies of *C*Las+ females. (C) Impacts of ds*DcDop2* treatment on mRNA levels of key genes in the AKH and JH signaling pathway in the ovaries of *C*Las+ females. (D-E) Rescue of the JH analogue and AKH on the fecundity and *C*Las titer phenotypes caused by the *DcDop2* knockdown. (F) JH titers in the abdomens of *C*las+ females after agomir-31a treatment for 48 h. (G) Effects of agomir-31a treatment on mRNA level of key genes in the AKH and JH signaling pathway in fat bodies of *C*Las+ females. (H) Effects of agomir-31a treatment on mRNA levels of key genes in the AKH and JH signaling pathway in the ovaries of *C*Las+ females. (I-J) Rescue of the JH analogue and AKH on the fecundity and *C*Las titer phenotypes caused by the agomir-31a treatments. Results in 6A-6C, 6E-6H, and 6J were displayed as means ± SEM with three independent biological replicates, each with three technical replicates. Data in 6D and 6I were shown as means ± SEM with thirty independent biological replicates. For figures D, E, I, and J, significant differences among the different treatments are indicated by lowercase letters based on one-way ANOVA followed by Tukey’s Honest Significant Difference tests at *p* < 0.05. Significant differences between treatments and controls are denoted by asterisks (Student’s *t*-test, \*\*\**p* < 0.001).

## Discussion

Plant pathogens manipulate the fitness of their vector insects to facilitate their own propagation and dissemination (Eigenbrode et al., 2018). Systemic infection by *C*Las induces alterations in the growth, development, behavior, metabolism, and reproduction of *D. citri* (Nian et al., 2024; Killiny and Jones, 2018; Hu et al., 2021). The substantial increase in the fecundity of *C*Las+ *D. citri* females undoubtedly exacerbates the occurrence and spread of HLB. Therefore, investigating the molecular mechanisms through which *C*Las enhances the reproductive capacity of *D. citri* can offer insights into the underlying mechanisms driving the rapid field transmission and prevalence of the pathogen, thereby providing novel perspectives for enhancing effective suppression of HLB. Dopamine, a pivotal neurotransmitter, can interact with other endocrine regulators to modulate the development and reproductive behavior of insect reproductive systems (Xu et al., 2017). Metabolomic analysis and ELISA measurements both indicate that *C*Las infection significantly elevates DA levels in *D. citri* females (Figure 1A-B). In various insects, DA levels have been closely linked to microbial infections. For example, bacterial and fungal infections have been shown to notably increase dopamine levels in the hemolymph of *Leptinotarsa decemlineata* (Say) (Coleoptera: Chrysomelidae) and *Galleria mellonella* (L.) (Lepidoptera: Pyralidae) (Chertkova et al., 2018). In *Bombyx mori* (L.) (Lepidoptera: Bombycidae), nucleopolyhedrovirus infection markedly boosts DA levels to enhance locomotor activity (Li et al., 2021). Therefore, further research is warranted to elucidate the role of DA in regulating the fecundity enhancement of *D. citri* induced by *C*Las infection.

In the DA biosynthesis pathway, tyrosine acts as the substrate for the production of an intermediate molecule, L-DOPA, which is then further converted to DA through the action of dopa decarboxylase (Siju et al., 2021). The synaptic release of DA is regulated by the vesicular amine transporter (VAT), which activates the DA receptor to initiate downstream signal transduction (Budnik and White, 1987). qRT-PCR analysis revealed a significant upregulation of four key genes (*DcHenna1*, *DcTh*, *DcDdc*, and *DcVat1*) involved in DA biosynthesis and release in *D. citri* females following *C*Las infection (Figure 1D). Subsequent knockdown of these four key genes resulted in a notable decrease in energy reserves, including TAG contents, glycogen abundance, and lipid droplets, along with reduced fecundity and *C*Las levels (Figure 2). These findings provide initial evidence that *C*Las manipulates DA biosynthesis to modulate its role in the enhanced fecundity of *D. citri* and proliferation of *C*Las within the psyllid. In *Solenopsis invicta* Buren (Hymenoptera: Formicidae), isolated, virgin females fed with a tyrosine hydroxylase inhibitor mixed in sucrose for 15 days showed significantly reduced egg laying and fewer chorionated oocytes in their ovarioles compared to females fed with sucrose alone. Restoring DA biosynthesis by supplementing the diet with L-DOPA reversed the effects on oogenesis and oviposition (Boulay et al., 2021). Furthermore, treatment with dsRNA or inhibitors targeting *DcDop2* expression inhibited DA signaling, resulting in similar phenotypic outcomes: decreased energy reserves, reduced fecundity, and *C*Las levels (Figure 3). In addition to DA receptors being implicated in behavioral phase transitions, conspecific olfactory attraction, and mating-related behaviors (Ma et al., 2011; Morigami and Sasaki, 2024; Guo et al., 2018), our results provide the first evidence for *DcDop2* involvement in the interaction between *C*Las proliferation and ovary development of *D. citri*.

In recent years, there has been growing research on miRNA regulation of the insect DA signaling pathway. For instance, miRNA-133 targets two key genes, *henna* and *pale*, in the pathway, downregulating their expression thereby inhibiting DA synthesis and inducing the transition of *L. migratoria* from gregarious to solitary behavior (Yang et al., 2014). The miRNA, let-7, enhances honey bee sensitivity to sucrose by targeting the DA receptor *AmDop2* (Liu et al., 2023). Our previous studies have demonstrated that miR-275 targets *DcVgR* and miR-34 targets *DcAKHR*, contributing to the enhanced fecundity in female *D. citri* induced by *C*Las (Nian et al., 2024; Li et al., 2024); this study has investigated whether *DcDop2* is post-transcriptionally regulated by miRNA and whether it is involved in the interaction between *D. citri* and *C*Las. Based on *in vivo* and *in vitro* experiments, we have demonstrated that miR-31a specifically binds to the 3’-UTR of *DcDop2*, negatively regulating its expression (Figure 4). The expression of miR-31a was significantly reduced in female *D. citri* 5 and 9 d after emergence (DAE) following *C*Las infection. Subsequent agomir-31a treatments resulted in a significant decrease in energy reserves, including TAG contents, glycogen levels, and lipid droplets, as well as reduced fecundity and *C*Las abundance; this mirrors the phenotypic effects of *DcDop2* interference (Figure 5). Therefore, we suggest that miR-31a negatively regulates *DcDop2* expression to modulate the fecundity enhancement of *C*Las-infected *D. citri* females and *C*Las proliferation. Further elucidation is needed on the regulatory interplay between miR-31a and other reported miRNAs.

In insects, the endocrine system plays a crucial role in coordinating the mechanisms governing energy metabolism and ensuring successful reproduction (Leyria, 2024). Individual neuroendocrine hormones not only regulate their specific target tissues but also engage in vital interactions among themselves, ensuring the successful progression of vitellogenesis and oogenesis. JH synthesis is stimulated by DA through the activation of the DA D1-like receptor in *D. melanogaster* (Rauschenbach et al., 2011). Following *DcDop2* knockdown or agomir-31a treatment, the JH levels in *C*Las-infected female *D. citri* were significantly reduced compared to the controls, indicating an influence of *DcDop2* on JH titers. Furthermore, qRT-PCR revealed a notable decrease in the mRNA expression of the JH receptor (*DcMet*), the key downstream gene, *DcKr-h1*, and three reproductive-related genes (*DcVg1-like*, *DcVgA1-like* and *DcVgR*) following *DcDop2* knockdown or agomir-31a treatment, as compared to the controls (Figure 6). In *D. melanogaster*, DA influences AKH secretion, and knockdown of the DA receptor in AKH cells results in starvation phenotypes (Braco et al., 2022). In our study, following *DcDop2* knockdown or agomir-31a treatment, the expression of *DcAKH* and *DcAKHR* exhibited a significant decrease compared to the controls (Figure 6). These findings suggest that DA signaling acts upstream of AKH and JH in the regulation of fecundity enhancement in *C*Las+ *D. citri* and *C*Las proliferation.

In summary, this study has revealed a new role of DA and its receptor in modulating the interaction between *C*Las and *D. citri*, extending beyond their known functions in development and behavioral transitions. As depicted in the conceptual model in Figure 7, *C*Las infection initially triggers the upregulation of DA biosynthesis genes, leading to increased DA levels. Subsequently, DA activates its receptor *DcDop2*, acting as an upstream regulator of AKH/JH signaling, governing energy metabolism and reproductive development. To maintain the appropriate level of *DcDop2* gene expression, miR-31a exerts negative regulation by targeting the 3’-UTR. Ultimately, *C*Las infection enhances the host’s reproductive capacity and promotes its own proliferation and dissemination, thereby achieving a win-win situation for both. Future investigations will address the following questions: (1) How does *C*Las manipulate the hormonal signals of citrus to enhance nutritional resources for *D. citri*, thus promoting psyllid fecundity and its own proliferation? (2) How do plant-derived cross-border miRNAs participate in regulating the interaction between *C*Las proliferation and development of *D. citri*?

**Figure 7.**
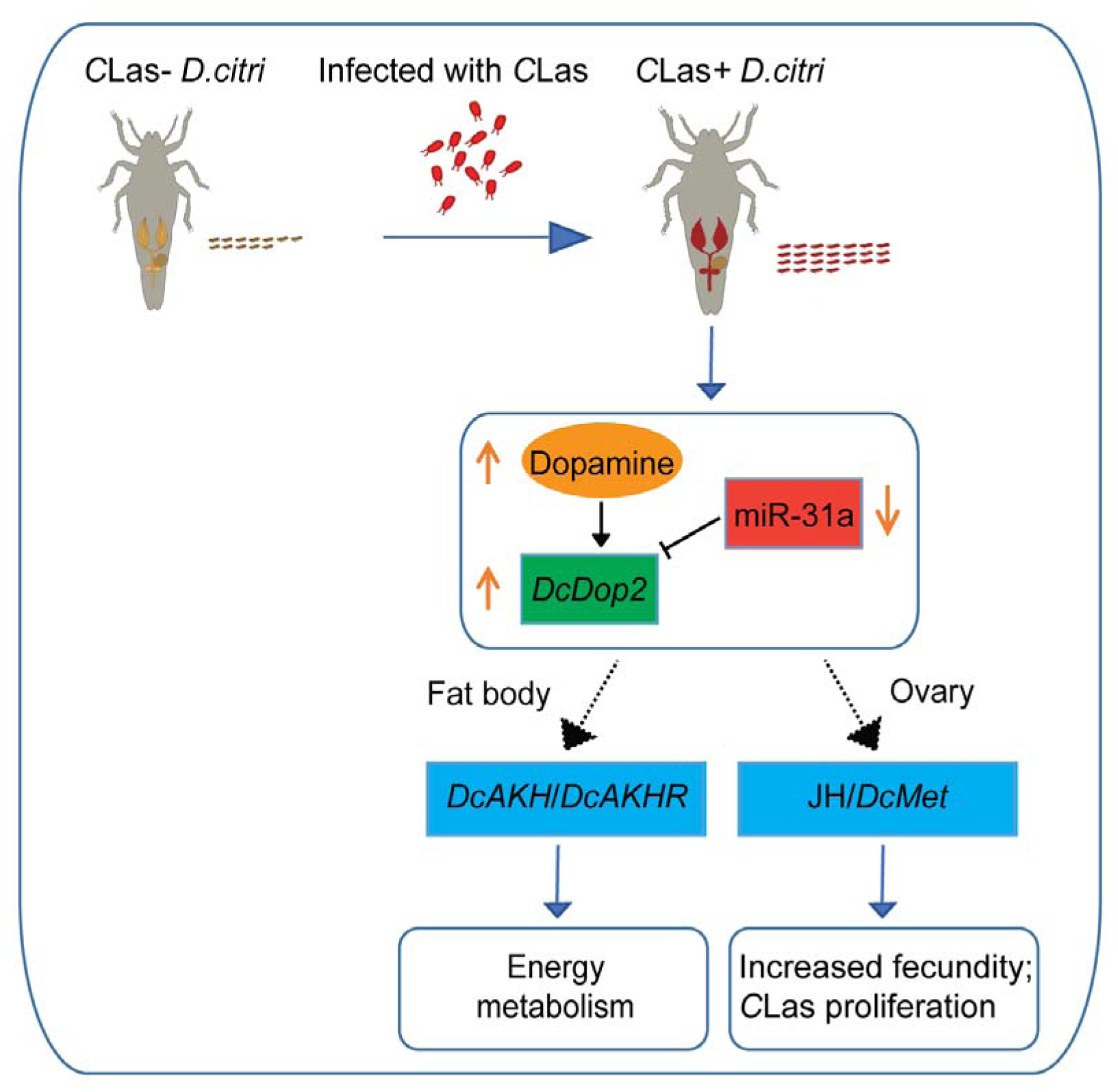
Diagram illustrating the involvement of DA and its receptor, *DcDop2,* in the coevolution between CLas and *D. citri*. *C*Las infection initially triggers the upregulation of DA biosynthesis genes, leading to increased dopamine levels. Subsequently, DA activates its receptor *DcDop2*, acting as an upstream regulator of AKH/JH signaling, governing energy metabolism and reproductive development. In order to maintain the appropriate level of *DcDop2* expression, miR-31a exerts negative regulation by targeting the 3’-UTR. Ultimately, *C*Las infection enhances the host’s energy metabolism, thereby boosting the host’s reproductive capacity and promoting its own proliferation and dissemination, achieving a win-win situation for both.

## Materials and Methods

### Host plants and insect populations

The *D. citri* colonies, both *C*Las-negative (healthy, *C*Las-) and *C*Las-positive (*C*Las+), were obtained from a laboratory cultures maintained continuously on healthy lemon (*Citrus* ×*limon* (L.) Osbeck) plants and *C*Las-infected lemon plants, respectively. Monthly monitoring of *C*Las infection in both the lemon plants and psyllids was conducted using quantitative polymerase chain reaction (qPCR) by the primers of *C*Las 16S ribosomal RNA gene (GenBank: L22532.1). The two populations were reared in separate incubators under similar conditions (temperature of 26 ± 1 LJ, relative humidity of 65 ± 5%, and a light: dark cycle of 14: 10). *C*Las-negative and *C*Las-positive lemon plants were cultivated in different glasshouses.

### Targeted metabolomic analysis

To assess the neurotransmitter contents in *C*Las+ *D. citri*, targeted metabolomic analyses were performed at Novogene Bioinformatics Technology Co. Ltd, (Beijing, China). In brief, a total of six samples, comprising three replicates each of *C*Las− or *C*Las+ females at 7-9 DAE, were prepared. The samples were firstly resuspended with liquid nitrogen and then diluted in water by vortexing. Subsequently, an ultra-high performance liquid chromatography coupled with tandem mass spectrometry (UHPLC-MS/MS) system (ExionLC™ AD UHPLC-QTRAP 6500+, AB SCIEX Corp., Boston, MA, USA) was used to quantify 23 targeted neurotransmitters. The mass spectrometer operated in negative multiple reaction mode (MRM) mode, with specific parameters set as follows: ionspray voltage (−4500 v), curtain gas (35 psi), ion source temp (550 LJ), ion source gas of 1 and 2 (60 psi). Differential metabolite screening was primarily based on Fold Change and P-value criteria, with the threshold set to FC > 1.1 or FC < 0.909 and *P* < 0.05 for identifying significant metabolites.

### Phylogenetic analyses

The physicochemical properties of *DcDop1* (GenBank accession number: XM_017446452.2), *DcDop2* (GenBank accession number: XM_026822145.1) and *DcDop3* (GenBank accession number: XM_008472271.3) were analyzed using the ProtParam tool available online (http://web.expasy.org/protparam/). Amino acid sequences of other insect species were retrieved and downloaded from the NCBI database. Phylogenetic trees were constructed using the neighbor-joining method in MEGA5.10 software, with 1000 bootstrap replicates. Bootstrap values exceeding 50% were displayed on the tree.

### Quantitative RT-PCR (qRT-PCR) for mRNA and miRNA

To analyze mRNA expression, total RNA was extracted using TRIzol reagent (Invitrogen, Carlsbad, CA, United States). First-strand cDNA synthesis was carried out with the PrimeScript™ II 1st Strand cDNA Synthesis Kit (Takara, Beijing, China) following the manufacturer’s instructions. qRT-PCR was performed using the TB Green® Premix Ex Taq™ II (Takara, Beijing, China) on an ABI PRISM® 7500 Real-Time System (Applied Biosystems, Foster City, CA, USA). The beta-actin (*Dc*β*-ACT*, GenBank XM_026823249.1) gene was used as the internal control for normalizing gene expression levels. For miRNA expression analysis, miRNA was extracted, synthesized, and quantified using the miRcute miRNA Isolation Kit, the miRcute Plus miRNA First-Strand cDNA Kit, and the miRcute Plus miRNA qPCR Kit (SYBR Green), all obtained from TIANGEN (Beijing, China). U6 snRNA was employed as an internal control to standardize miRNA expression levels. Detailed primer information is provided in Table S1. Three technical replicates for each sample were performed on the same plate. Each experiment was repeated three times. The reaction with RNase-free water as the template was designed as the negative control.

### Measurement of TAG levels

To determine TAG levels, a Triglycerides Colorimetric Assay (Cayman) was employed following the manufacturer’s instructions. In brief, thirty fat bodies were homogenized in 100 μL of Diluent Assay Reagent. Subsequently, 10 μL of the supernatant was incubated with Enzyme Mixture solution, and the resulting TAG contents were standardized to the protein levels in the supernatant, measured using a BCA protein assay (Thermo Fisher Scientific). Each treatment underwent analysis with three independent biological replicates, each consisting of three technical replicates.

### Measurement of glycogen levels

Glycogen levels were determined using a Glycogen Assay Kit (Cayman) according to the manufacturer’s instructions. Thirty fat bodies were homogenized in 100 μL of Diluent Assay Reagent, and 10 μL of the supernatant was then incubated with Enzyme Mixture solution and Developer Mixture. The measured glycogen levels were normalized to the protein levels in the supernatant and quantified using a BCA protein assay (Thermo Fisher Scientific). Each treatment was assessed with three independent biological replicates, each comprising three technical replicates.

### Nile red staining

Dissected fat bodies were fixed in 4% paraformaldehyde for 30 min at 25 °C, followed by two washes with 1 × PBS. Lipid droplets were incubated for 30 min in a mixture of Nile red (0.1 μg/μL, Beijing Coolaber Technology Co., Ltd) and DAPI (0.05 μg/μL), then washed twice with 1 × PBS. Imaging of the samples was performed using laser scanning microscopy (TCS-SP8, Leica Microsystems Exton, PA, USA). Each experiment was repeated three times.

### Assessment of reproductive parameters

Treated females from the two colonies were paired with healthy males for testing. A single female and male were placed onto young vegetative shoots (flush) of healthy lemon plants to induce oviposition. The flush and insects were enclosed in tied, white, mesh bags (150 × 200 mm). These lemon plants were kept in an incubator under conditions of 25 ± 1 °C, 65 ± 5% RH, 14L: 10D photoperiod. After 24 h, the preoviposition period and oviposition period were recorded, the number of eggs laid per female were counted, and the psyllid pair was transferred to new flushes for continued egg laying. Egg counts were conducted daily until all females perished. All experiments were replicated three times, with approximately 15 pairs of *D. citri* used for each replication.

### Double-stranded RNA synthesis and RNAi experiments

Double-stranded RNAs (dsRNA) targeting *DcHenna1*, *DcTh*, *DcDdc*, *DcVat1*, and *DcDop2* were synthesized as the kit’s instructions using sequence-specific primers (Table S1). The dsRNA was synthesized by using a Transcript Aid T7 High Yield kit (Thermo Scientific, Wilmington, DE, United states) and purified with the GeneJET RNA Purification kit (Thermo Scientific). RNAi treatments were administered by feeding dsRNA via an artificial diet, following established protocols (Nian et al., 2024). Briefly, a glass cylinder (25 × 75 mm) was used as the feeding chamber. Fifteen females at 7 DAE (for dsRNA and miRNA agomir) from *C*Las-negative and *C*Las-positive *D. citri* were fed with an artificial diet (200 μL) placed between two layers of stretched parafilm. The artificial diet consisted of 20% (w: v) sucrose mixed with dsRNA (250 ng/μL) or miRNA antagomir/agomir (7.5 μM). The ovary samples were collected at 48 h following dsRNA/miRNA antagomir/agomir feeding for qRT-PCR and western blot analysis to confirm the interference efficiency. For the validation assays of miRNA/genes involvement in reproductive regulation after feeding with dsRNA/miRNA agomir for 48 h, the treated females are divided and used for three assays. The first assay was conducted to investigate the reproductive parameters (the pre-oviposition period, the oviposition period and fecundity) following the above methods. The second assay was carried out to detect *C*Las titer in the ovary. The third assay was performed to observe the ovary morphology. Feeding ds*GFP* was used as the control when psyllids were fed with dsRNA. Antagomir/agomir negative control was used as the control when psyllids were fed with miR-31a antagomir/agomir. All the experiments were performed across three separate time replicates, with 20 pairs of *D. citri* for each. Photographs of ovaries were captured by an Ultra-Depth Three-Dimensional Microscope (VHX-500). Fat bodies were stained with Nile red, TAG levels, and glycogen levels in the fat bodies were evaluated. For the rescue experiments, the females were first fed with ds*DcDop2* for 48 h, then the relative neuropeptide hormone (DA, JHA, or AKH) was treated. The corresponding parameters were analyzed based on the above methods.

### Luciferase activity assay

To investigate the regulatory effects of miR-31a, miR-275, and novel-miR-15 on the 3’-UTR of *DcDop2*, a 147 bp sequence surrounding the predicted target sites was cloned into the pmirGLO vector (Promega, Wisconsin, USA) to generate the *DcDop2*-3’UTR-pmirGLO plasmid, utilizing the pEASY®-Basic Seamless Cloning and Assembly Kit. A mutated version, *DcDop2*-3’UTR mutant-pmirGLO plasmid, was also created. Following standard protocols, 500 ng of the constructed vectors (*DcDop2*-3’UTR-pmirGLO plasmid or *DcDop2*-3’UTR mutant-pmirGLO plasmid) and 275 nM of miRNA agomir/agomir-NC were co-transfected into HEK293T cells using a Calcium Phosphate Cell Transfection Kit (Beyotime, Nanjing, China). After 24 h of co-transfection, the activities of firefly and Renilla luciferase were measured using the Dual-Glo Luciferase Assay System (Promega, Madison, WI, USA) with a plate reader.

### Fluorescence in situ hybridization (FISH)

For FISH analysis using a *C*Las probe, ovaries were fixed in Carnoy’s fixative (glacial acetic acid-ethanol-chloroform, 1: 3: 6, vol/vol/vol) for 12 h at 25 _. The fixed tissues underwent several washes and pre-incubation steps before hybridization with the probe. Subsequently, the samples were stained with DAPI and mounted for visualization using a Leica TCS-SP8 confocal microscope. Excitation lasers at specific wavelengths were used to detect different signals, and image processing was conducted with Leica LAS-AF software. Control experiments without the probe and CLas-negative controls were included for specificity verification. Three FISH tests were performed, with a minimum of 15 ovaries examined in each test for consistency. The probe sequences are detailed in Table S1.

### Determination of JH titers and Dopamine levels

JH titers were quantified using the Insect JH ELISA kit (ml077240; Shanghai Enzyme-linked Biotechnology Co., Ltd. Shanghai, China) according to the manufacturer’s protocol (Nian et al., 2024). Briefly, 50 µL of the homogenized tissue sample was added to each well. Subsequently, 100 µL of HRP-conjugated detection antibody was added. After a series of standard incubation, washing, and stop procedures, the OD value of each well at 450 nm was measured by a Molecular Devices i3x microplate reader. Dopamine levels were assessed with a DA ELISA kit (ml077133; Shanghai Enzyme-linked Biotechnology Co., Ltd. Shanghai, China). Tissue samples were homogenized, and the supernatants were incubated with the respective antibodies. Dopamine contents were normalized to standard sample OD values determined by a Molecular Devices i3x microplate reader. Each treatment was analyzed with three independent biological replicates, each including three technical replicates.

### Western blotting

*C*Las-positive females at 7 DAE, treated with ds*DcDop2* and miR-31a agomir for 48 h during ovarian development, were collected as described above. Total proteins were extracted from the dissected ovaries of the psyllids using RIPA protein lysis buffer (50 mM Tris pH 7.4, 150 mM NaCl, 1% Triton X-100, 1% sodium deoxycholate and 0.1% SDS) supplemented with 1 mM PMSF. The lysates were then cleared by centrifugation at 4 °C for 20 minutes. Extracted proteins were quantified using a BCA protein assay kit (Beyotime, Jiangsu, China). Equal amounts of protein (50 μg) were separated by 12% SDS-PAGE and subsequently transferred to polyvinylidene fluoride membranes (Millipore). The membranes were blocked with 5% nonfat powdered milk to reduce non-specific binding and incubated overnight at 4 °C with the primary antibody (*DcDop2*; 1: 1000 dilution, ABclonal Technology Co., Ltd., Wuhan, China). Afterward, the membranes were incubated for 2 hours at 25 °C with the secondary antibody (goat anti-rabbit IgG conjugated with HRP, 1: 10000 dilution). A mouse monoclonal antibody against β-actin (TransGen Biotech, Beijing, China) was used as a control. Immunoreactivity was visualized using enhanced chemiluminescence with the Azure C600 multifunctional molecular imaging system (USA). Each experiment was repeated two times.

### RNA immunoprecipitation (RIP)

A RIP experiment was conducted using a Magna RIP kit (Millipore, Billerica, MA) according to the manufacturer’s instructions. Females at 7 DAE were fed with a miR-31a agomir for 12 hours and then subjected to RIP analysis 10 hours later. The ovaries were dissected and homogenized in ice-cold RIP lysis buffer. The homogenates were stored at −80 °C overnight. A total of 5 μg of Ago-1 antibody or normal mouse lgG (Millipore), which served as a negative control, was pre-incubated with magnetic beads. The frozen homogenates in the RIP lysates were thawed and centrifuged, and the supernatants were incubated with the magnetic bead-antibody complex at 4 °C overnight. 1/10 of the lysate was reserved as an “input” sample. The immunoprecipitated RNAs were reverse transcribed into cDNA using random hexamers, and qRT-PCR was performed to quantify miR-31a and *DcDop2*. The “input” samples and lgG controls were analyzed to normalize the relative expression levels of the target gene. Each experiment was repeated three times.

### Heterologous expression and calcium mobilization assay

The full ORF of *DcDop2* was successfully cloned into the expression vector, pcDNA3.1, and the correctness of the cloning was verified through sequencing performed by TSINGKE Bio. Endotoxin-free plasmid DNA was extracted from the vector, and dopamine was procured from MedChemExpress (HY-B0451A). Chinese hamster ovary (CHO-WTA11) cells engineered with Gα16 subunit and aequorin, were utilized for the experiments. These cells were transfected with either pcDNA3.1-*DcDop2* or empty pcDNA3.1 vector (negative control) using Lipofectamine 2000 (Thermo Fisher Scientifc, Waltham, MA USA). Calcium mobilization assays were conducted with minor adjustments following a previously outlined protocol (Zhang et al., 2023). Briefly, 48 h post-transfection, the cells were incubated in the dark with coelenterazine h (Promega, Madison, WI, USA) for 3 h. Subsequently, various concentrations of dopamine (10^−7^, 10^−6^, 10^−5^, 10^−4^, 10^−3^, 10^−2^, 10^-1^, and 1 mM) were added to opaque 96-well plates. Luminescence was measured for 15 s using a SpectraMax i3x Multi Mode Microplate Reader (Molecular Devices). Each experiment was repeated three times.

### Statistical analysis

Statistical analyses were conducted using GraphPad Prism 8.0 software. All data are presented as means ± standard errors of mean (SEM). Pairwise comparisons were assessed using Student’s *t*-tests, with significance levels denoted as follows: \**p* < 0.05, \*\**p* < 0.01, \*\*\**p* < 0.001. For multiple comparisons, one-way ANOVA coupled with Tukey’s Honest Significant Difference tests was employed to determine significant differences at *p* < 0.05.

## Acknowledgments

This research was supported by the National Natural Science Foundation of China (32572827), Guangdong Academy of Agricultural Sciences Special Fund for Talent Recruitment in Science and Technology (R2025YJ-QG002), and the Open Competition Program of Ten Major Directions of Agricultural Science and Technology Innovation for the 14th Five-Year Plan of Guangdong Province (2022SDZG07).

## Author contributions

S.Z. and X.N. designed research; S.Z., J.L., and X.N. performed research; J.H., W.Y. P.H., G.A.C.B., Y.C., and Y.H. contributed new analytic tools; S.Z., and X.N. analyzed data; S.Z., X.N., P.H., and G.A.C.B. wrote the paper; X.N., and Y.H. provided the fund.

## Benefit-sharing statement

The authors declare no conflict of interest.

## Data availability statement

All data are available in the manuscript or the supplementary materials.

## Supplementary data

**Figure S1.**
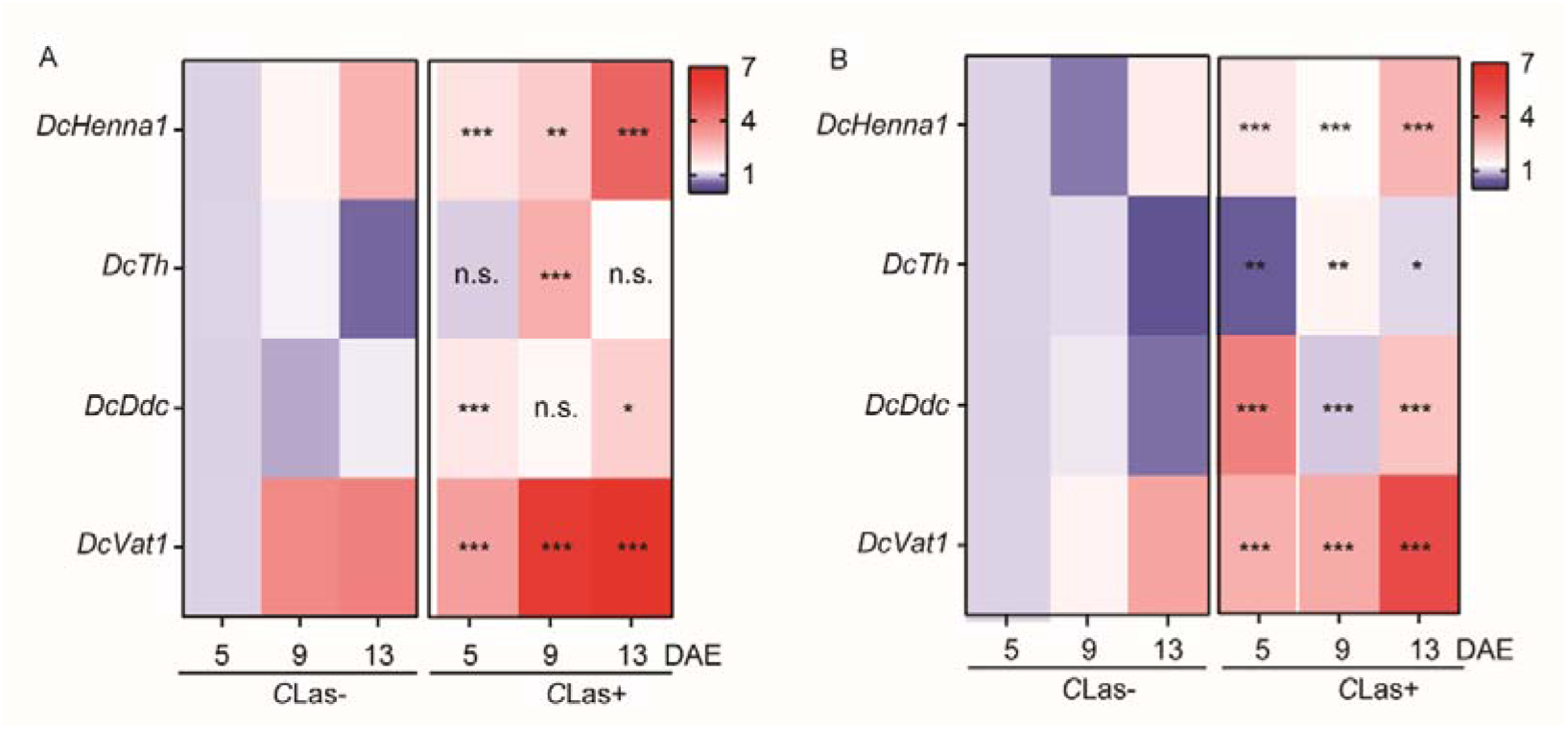
The expression levels of *DcHenna1*, *DcTh*, *DcDdc*, and *DcVat1* in the whole body and head of *C*Las− and *C*Las+ psyllids. Data are presented as means ± SEMs. Significant differences between *C*Las− and *C*Las+ psyllids were determined using Student’s *t*-tests (\**p* <0.05, \*\**p* < 0.01, \*\*\**p* < 0.001).

**Figure S2.**
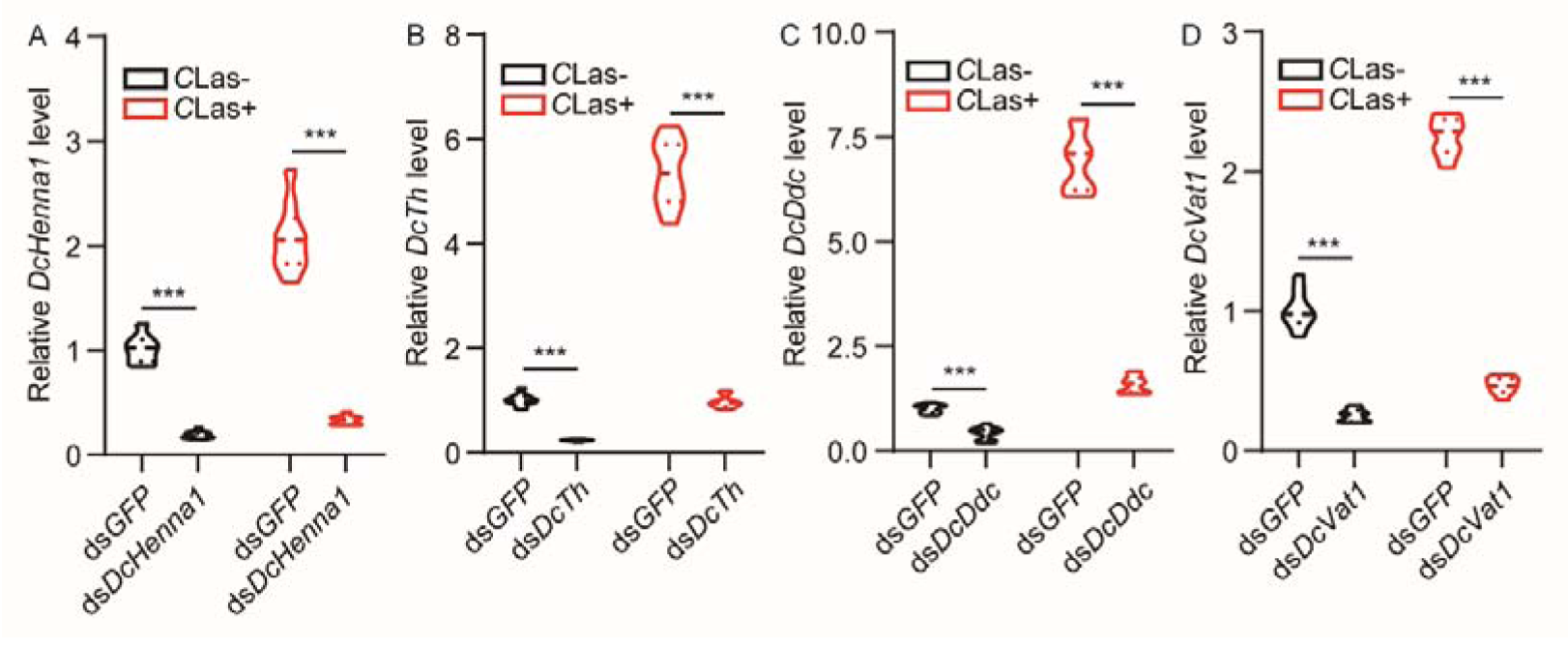
The RNAi efficiency of *DcHenna1*, *DcTh*, *DcDdc*, and *DcVat1* in *C*Las− and *C*Las+ psyllids treated with ds*RNA* for 48 h.

**Figure S3.**
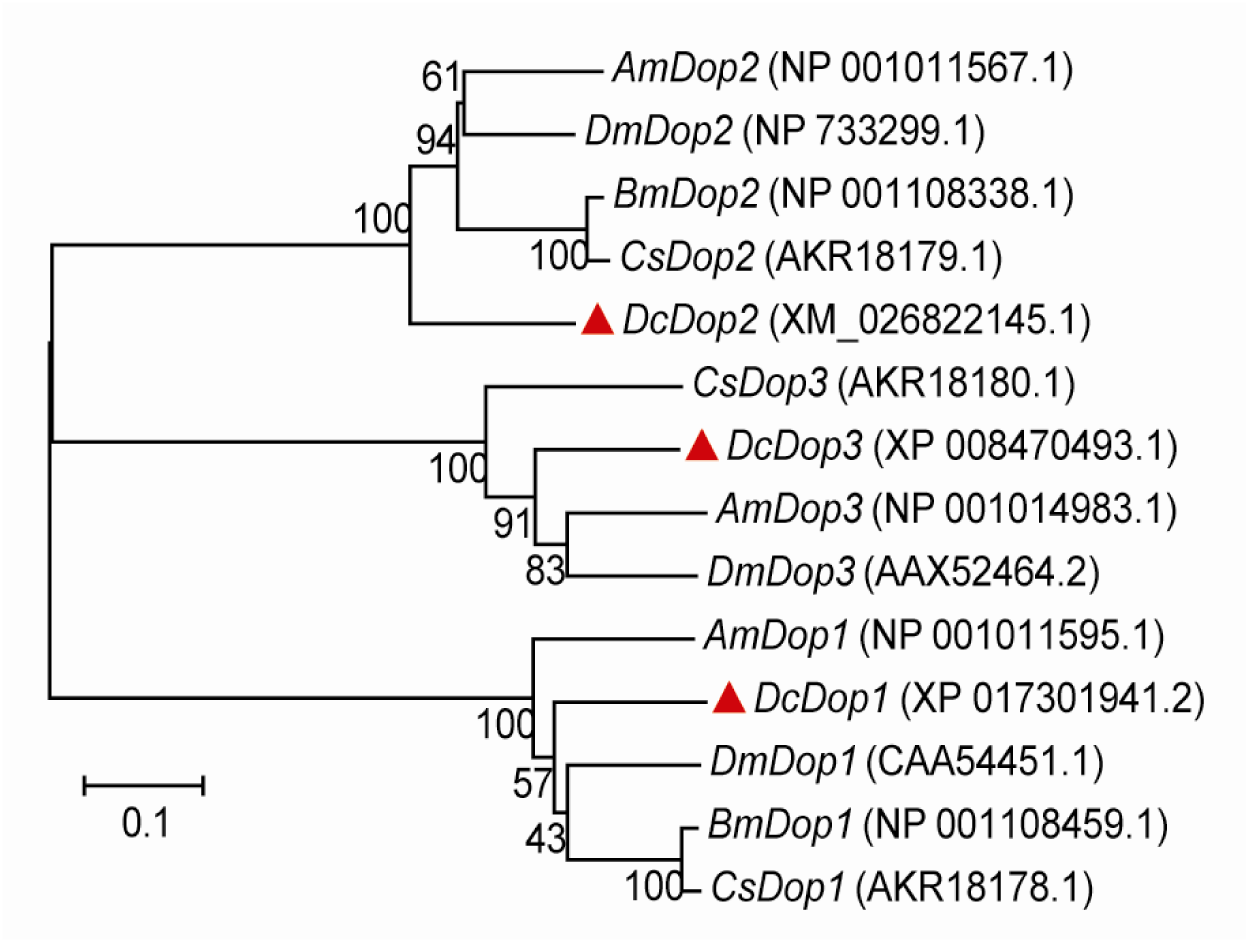
Phylogenetic analysis the protein sequences of *DcDop1*, *DcDop2*, and *DcDop3*. The phylogenetic tree was constructed using neighbor-joining with 1000 bootstrap replicates; values > 50% are shown on the tree. The scale bar indicates the number of amino acid substitutions per site. Dc = *Diaphorina citri*; Dm = *Drosophila melanogaster*; Bm = *Bombix mori*; Cs = *Chilo suppressalis*.

**Figure S4.**
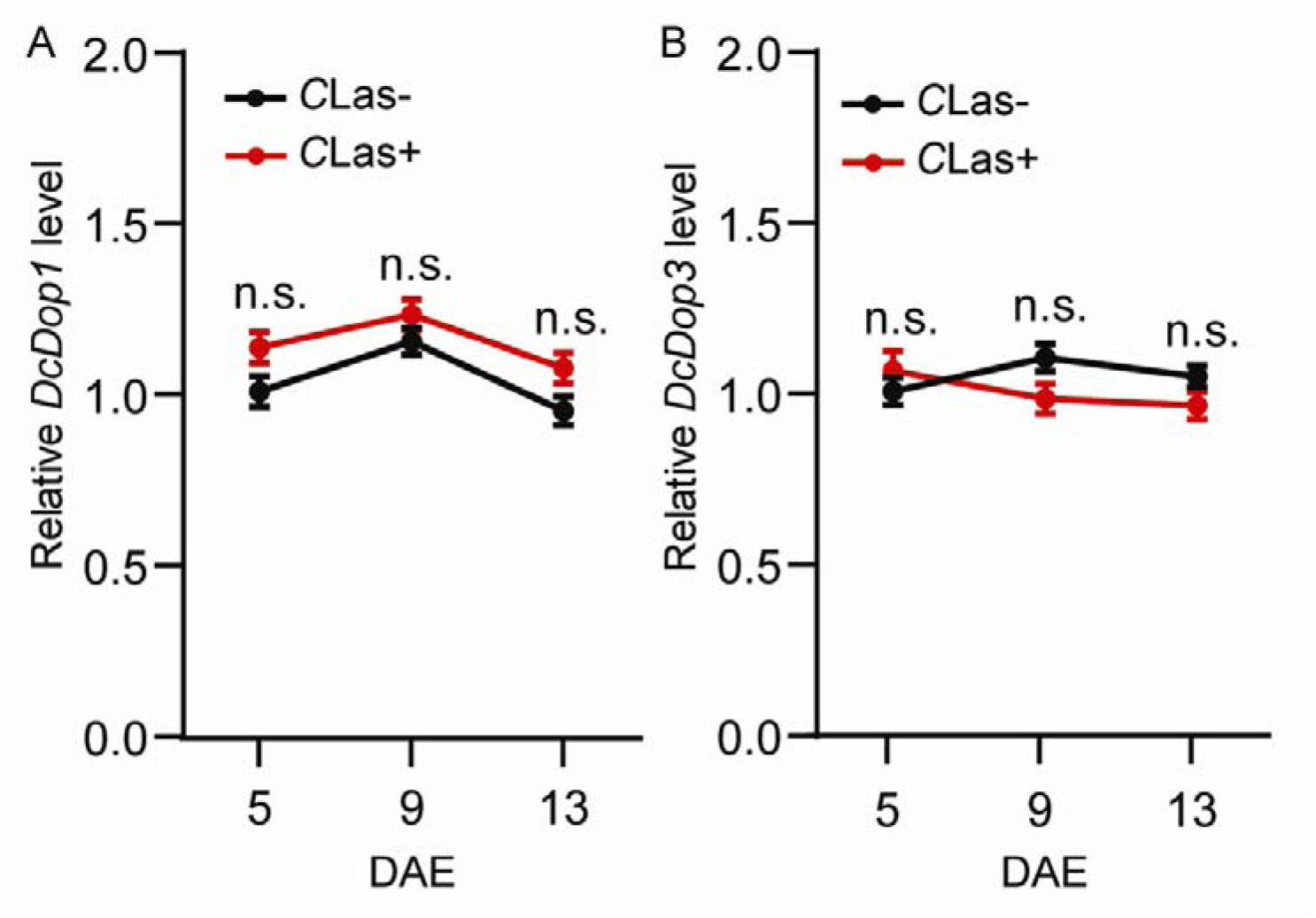
Temporal expression patterns of (A) *DcDop1* and (B) *DcDop3* in the ovaries of *C*Las− and *C*Las+ psyllids.

**Figure S5.**
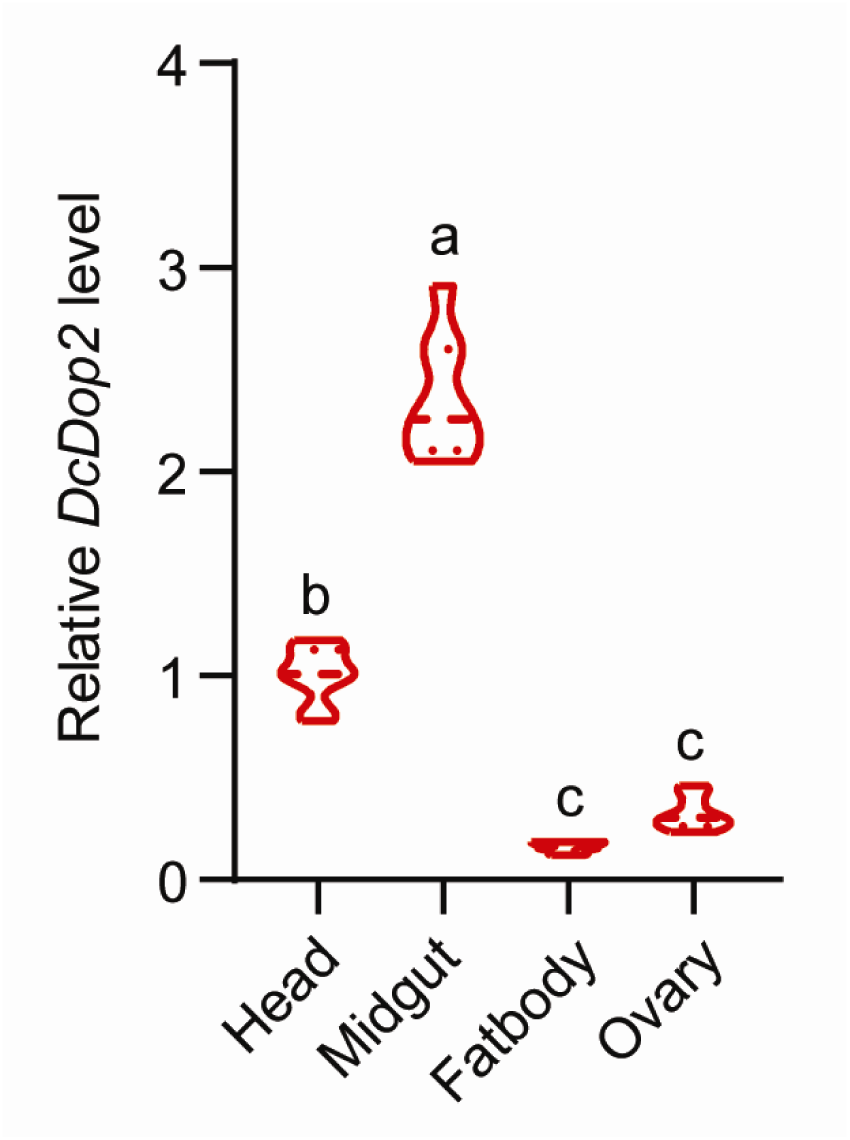
Expression of *DcDop2* in the head, midgut, fat body, and ovary of adult female *C*Las+ 7 DAE.

**Figure S6.**
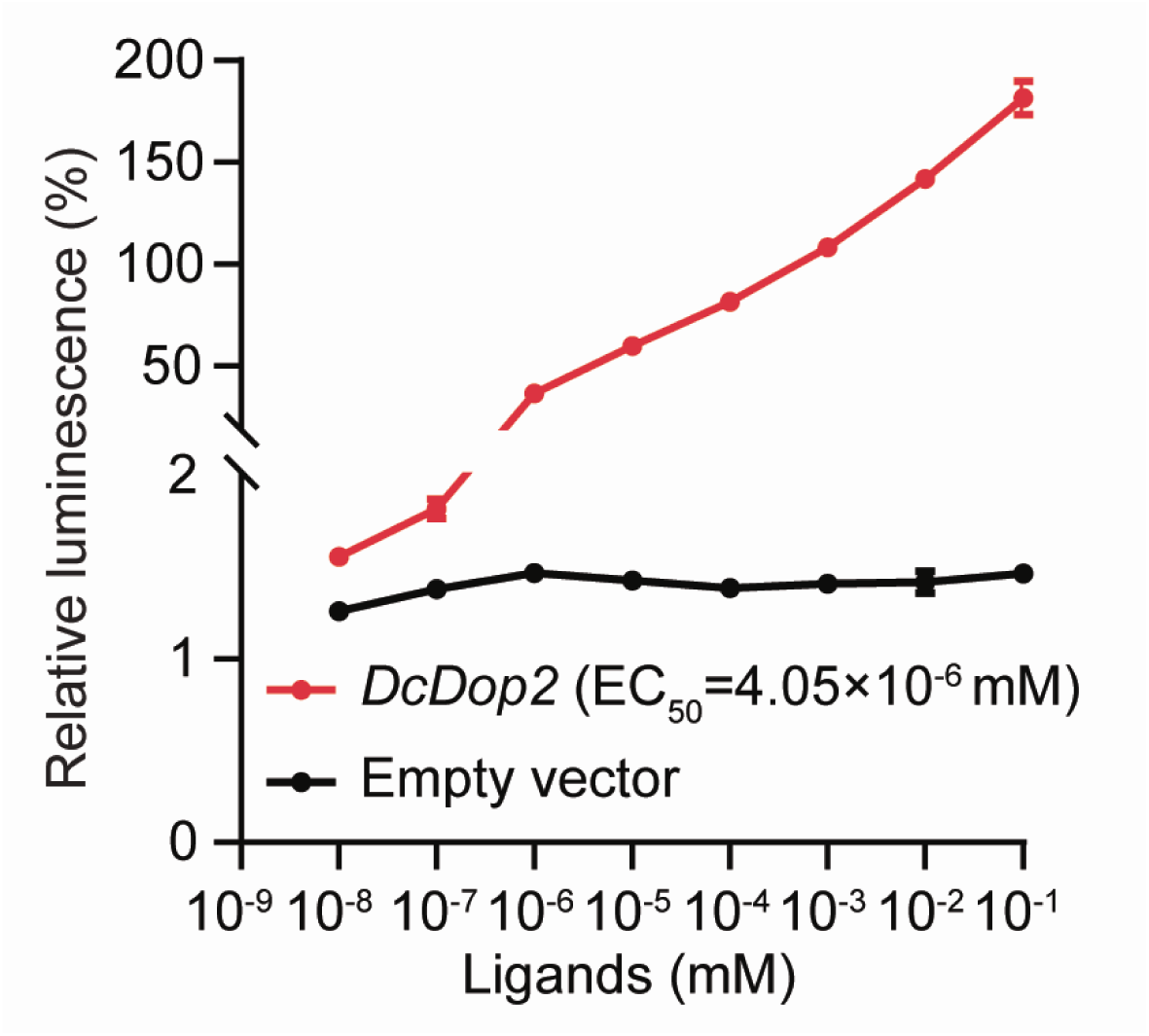
Concentration-response curves for the effect of Ca^2+^ on *DcDop2*-expression in CHO cells.

**Figure S7.**
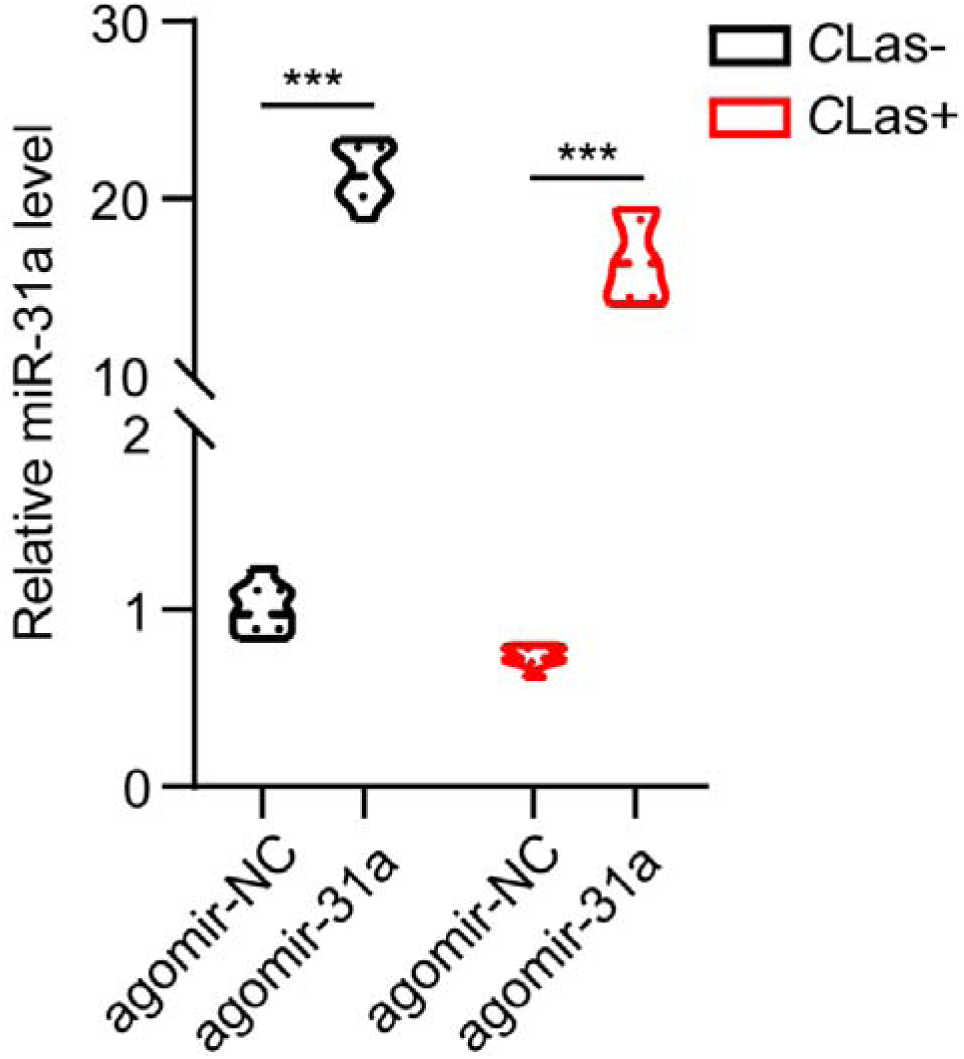
The expression levels of miR-31a in *C*Las-negative and *C*Las-positive females treated with agomir-31a or agomir-NC for 48 h.

**Table S1.**
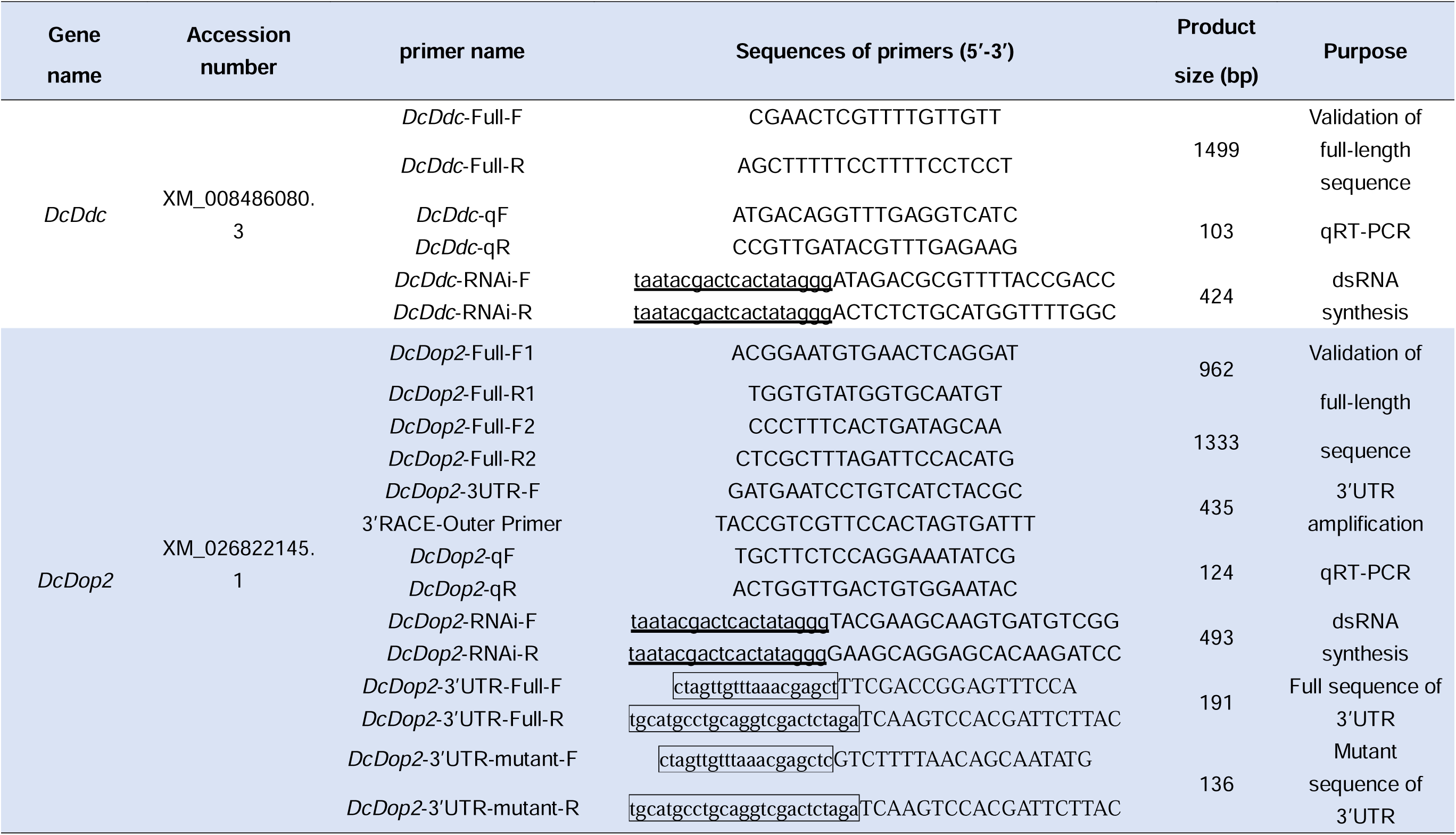
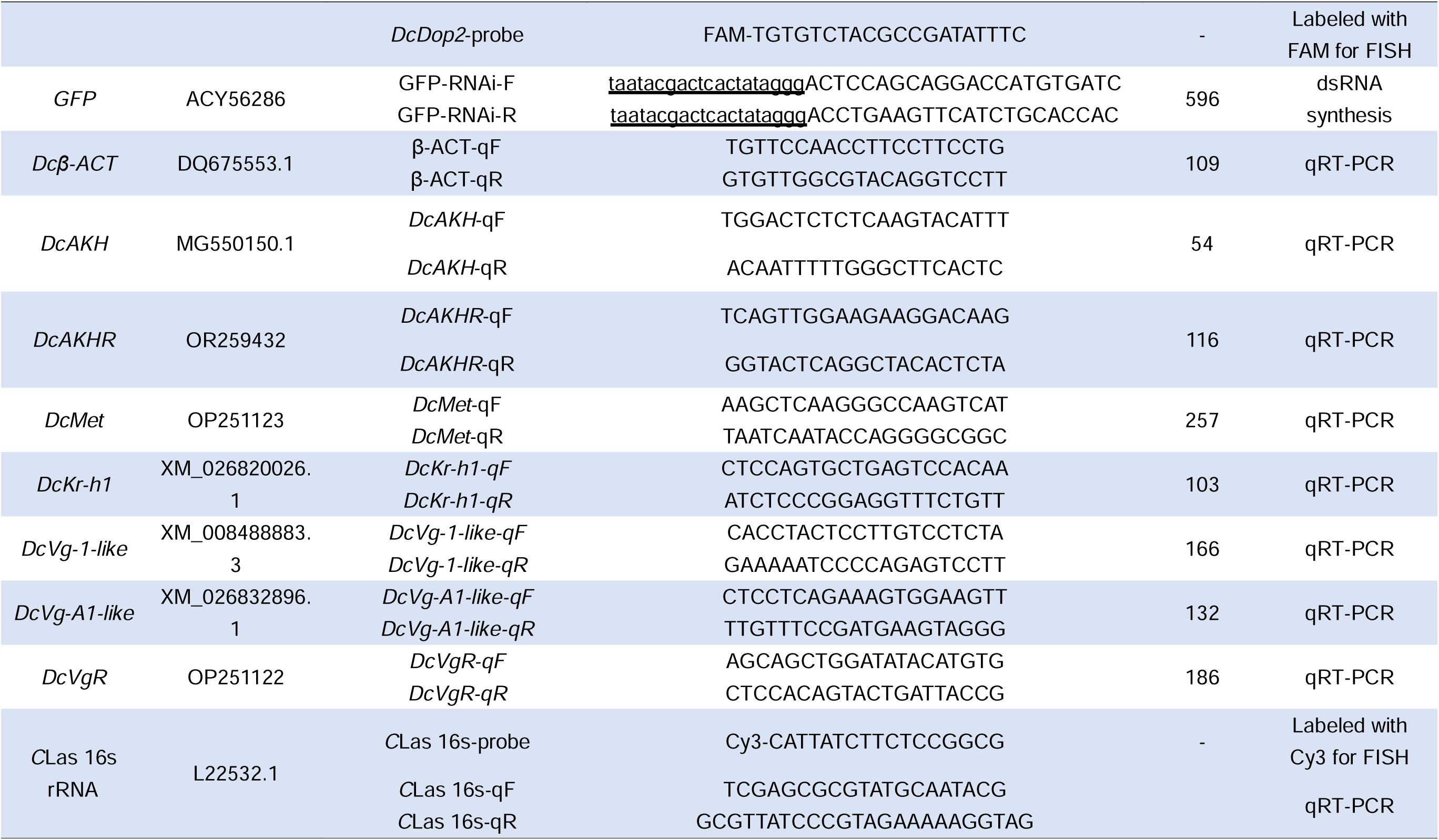
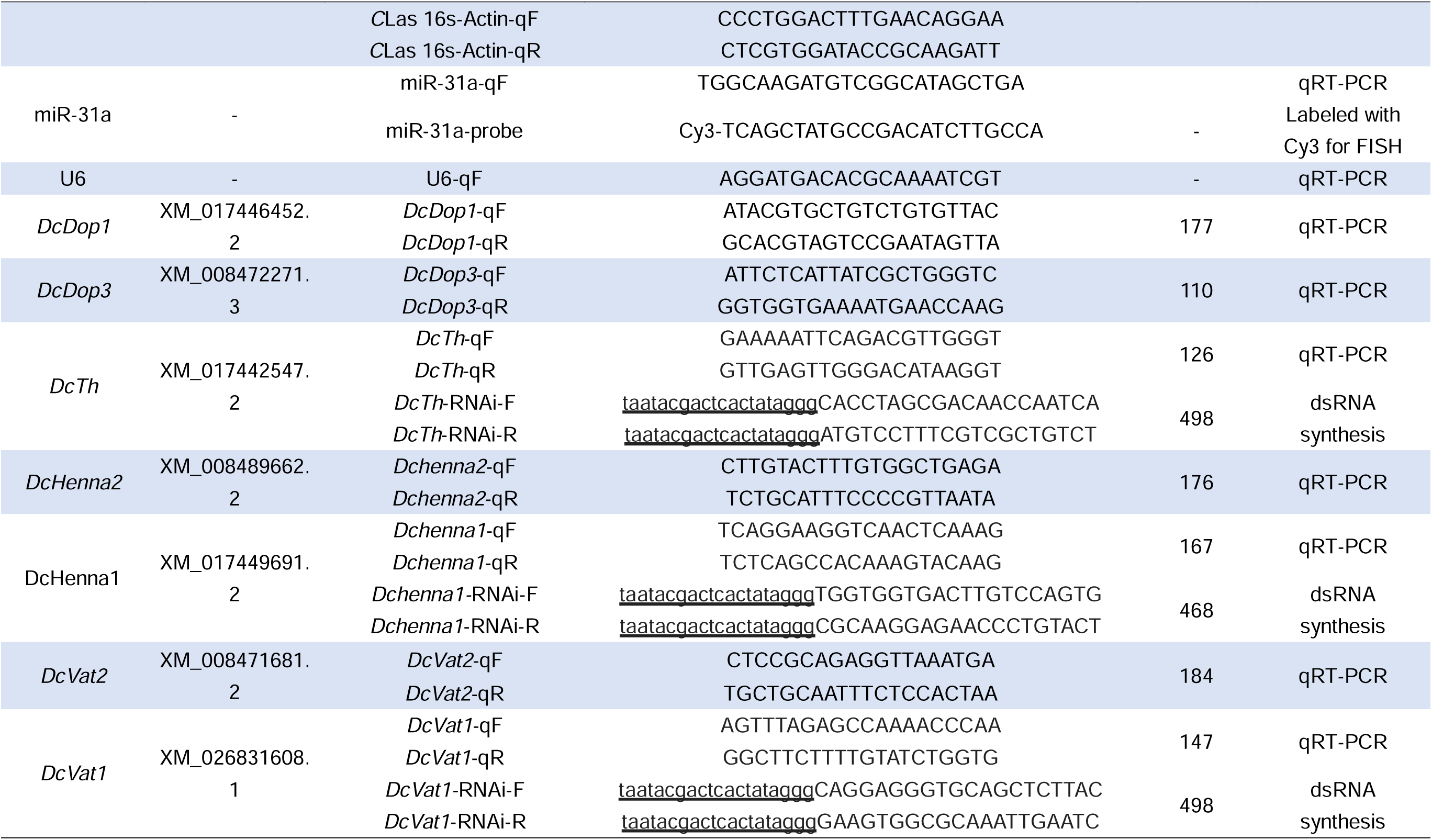
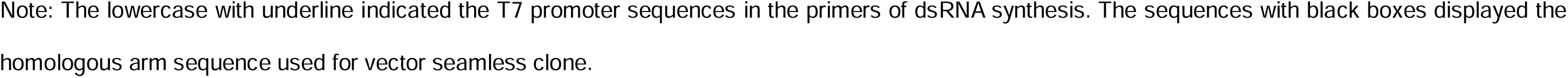
The primers used in this study.

